# Predicting Epitope Candidates for SARS-CoV-2

**DOI:** 10.1101/2022.02.09.479786

**Authors:** Akshay Agarwal, Kristen L. Beck, Sara Capponi, Mark Kunitomi, Gowri Nayar, Edward Seabolt, Gandhar Mahadeshwar, Simone Bianco, Vandana Mukherjee, James H. Kaufman

## Abstract

Epitopes are short amino acid sequences that define the antigen signature to which an antibody binds. In light of the current pandemic, epitope analysis and prediction is paramount to improving serological testing and developing vaccines. In this paper, we leverage known epitope sequences from SARS-CoV, SARS-CoV-2 and other Coronaviridae and use those known epitopes to identify additional antigen regions in 62k SARS-CoV-2 genomes. Additionally, we present epitope distribution across SARS-CoV-2 genomes, locate the most commonly found epitopes, discuss where epitopes are located on proteins, and how epitopes can be grouped into classes. We also discuss the mutation density of different regions on proteins using a big data approach. We find that there are many conserved epitopes between SARS-CoV-2 and SARS-CoV, with more diverse sequences found in Nucleoprotein and Spike Glycoprotein.

## 1 Introduction

Since the first case of COVID-19 was reported in December 2019, the prevalence of SARS-CoV-2 has grown at an incredible pace resulting in a worldwide pandemic. The scale and rate of spread of the disease continues to grow. As a consequence, humans have had to drastically change their lifestyle, and scientists and clinicians have been presented with great challenges such as identifying and studying rapidly emerging new strains and developing vaccines.

In order to assist with pandemic response and to further the understanding of the adaptive immune response to the virus, we decided to identify, and investigate the presence and evolution of SARS-CoV-2 epitopes, specifically for Nucleoprotein (N) and Spike glycoprotein (S). In this paper, we present our analysis of 11 different SARS-CoV-2 proteins amounting to 28K unique protein sequences. We use data from early on in the pandemic and show the relevance of results for Delta variant as well, thus underscoring the stability of top epitopes. We highlight in detail the results obtained for Nucleoprotein and Spike glycoprotein since these proteins are of particular medical interest^1^. Below, we give a short introduction of epitopes, Nucleoprotein and Spike glycoprotein.

Epitopes are peptides on the surface of an antigen to which an antibody binds. Each epitope is defined by a unique sequence of amino acids. With considerations regarding protein structure, epitopes are broadly classified into two categories: linear epitopes that are continuous amino acid sequences, and non-linear or conformational epitopes that are discontinuous peptides on the unfolded sequence. Non-linear epitopes may only be recognized by the immune system depending on neighboring loci. In this investigation, we focus on linear epitopes.

Further, based on function, epitopes are of two types: T-Cell epitopes and B-Cell epitopes. T-Cell epitopes bind to the major histocompatibility complex (MHC) molecules found on the cell surface. The corresponding antigens vary between 8-17 amino acids in length. They are recognized by CD8 and CD4 T-cells^2^. B-Cell epitopes are solvent-exposed portions of the antigen that bind to secreted and cell-bound immunoglobulins i.e. the ones to which antibodies bind^2^. Epitope-based vaccines contain isolated B-Cell or T-Cell epitopes and typically contain multiple distinct epitopes in order to increase the effectiveness of the vaccine given the time evolution of the virus.

The structural Nucleoprotein (N) is the main virion component. It encapsulates the negative-stranded RNA. The viral genomics RNA and the N protein assemble into the ribo-nucleoprotein which interacts with the membrane (M) protein and is packaged into virions^3^. N proteins are also involved in other viral functions including mRNA transcription, replication^4,5^ and immune regulation^6-8^. For other RNA viruses including influenza, N sequence is often used for species identification^9^. N proteins have two functionally distinct, conserved structural domains, the N-terminal RNA-binding domain (NTD) and the C-terminal domain (CTD) responsible for both RNA-binding and dimerization^10^, connected by an intrinsically disordered region (IDR) called linker and flanked by intrinsically disordered regions (IDRs). Due to the large disordered regions, the whole N protein structure has not been resolved yet, but the N-NTD and N-CTD domains have been solved at high resolution for different human-infecting Coronaviridae^11–15^, including SARS-CoV-2^16-19^.

The Spike glycoprotein (S) is important as it plays a vital role in virus binding, fusion and entry into the host cell^20–27^. The S protein is a homotrimeric class I fusion protein and each protomer comprises of two functional sub-units, S1 and S2. The S protein is found on the surface of the viral membrane in the metastable prefusion configuration and undergoes major conformational changes to allow membrane fusion with the host cell^21,22,24,26–29^. This process occurs after the receptor binding domain (RBD) in S1 binds to the host cell receptor. The RBD is flanked by the NTD and the CTD (CTD1, CTD2)^21^. There are two cleavage sites and one of them separates S1 from S2 fragment. During cell entry S1 dissociates and S2 refolds into a stable post-fusion state^30-32^. S2 comprises the fusion peptide (FP), the FP proximal region (FPPR), the heptad repeat 1 (HR1), the central helix (CH), the connector domain (CD), the heptad repeat 2 (HR2), a transmembrane domain (TM), and the cytoplasmic tail (CT). Comparison between pre- and post-fusion structures suggests that HR1 transitioning movements allow insertion of the FP into the host cell membrane and folding back of HR2^21^. In the post-fusion structure, HR1 and CH form a ≈ 180 Å extended, three-stranded coiled coil.

There were two primary motivations for studying and predicting epitopes in SARS-CoV-2 and leveraging known epitopes of other Coronaviridae to do so. First, this can help improve epidemiological models. Since the reservoir for most emerging infectious disease is the wild animal population, a virus newly infecting and spreading in human populations requires that the virus successfully hop between species, reproduce within the cells of the new host, and transmit between individuals with an effective reproductive number *R_t_*>1. The latter condition requires that a majority of the human host population be susceptible to the virus or the virus have a high *R*_0_. In general, the initial susceptible fraction is less than 100% (it must by greater than the inverse of the basic reproductive number, S/N > 1/*R*_0_). Understanding the initial susceptible fraction (S/N) is important to epidemiological models and forecasts. The existence of SARS-CoV-2 epitopes with identical sequence across known species may provide insight into the fraction of the population initially resistant to the virus based on an adaptive immune response developed from exposure to other pathogens. Second, the adaptive immune system of two individuals infected by the same organism may learn different peptide sequences and, therefore, produce antibodies for different epitopes. Conversely, since some protein domains are common between organisms, an adaptive immune response developed based on exposure to one organisms may sometimes increase risk against a different infectious pathogen^33^.

Identification of common epitopes may also help pharmaceutical research groups to identify therapeutics more quickly by looking at existing treatments for other organisms. Identifying conserved targets may also aid vaccine design and development.

In this investigation, across all proteins studied, we identified 112 lab-confirmed B-Cell epitopes of which 3 were found on SARS-CoV-2 while the rest were originally found in the SARS-CoV virus identified in 2003. In addition, 279 lab-confirmed T-Cell epitopes were identified of which 221 were found on SARS-CoV-2 itself while the rest were originally identified in SARS-CoV. Additionally, we predict 237 B-Cell and 110 T-Cell new epitopes based on fuzzy search techniques.

A newly emerging human virus will adapt to the environment presented by its new host’s by selection of mutations in some proteins, with other regions remaining conserved. In order to study these mutations and analyze if there is a correlation between them and the host a virus is found in, we analyze how the occurrence of different groups of epitopes vary by host. Given a common set of epitopes between SARS-CoV-2 and other Coronaviridae, we perform hierarchical clustering of the epitopes to identify related epitopes that share sequence (overlapping the same protein region). Prediction of previously unknown epitopes and identification of their location on specific proteins can indicate which epitopes are evolving rapidly and which may be more stable and better candidates for vaccine targets.

## 2 Data

Data for this study was obtained primarily from two sources. Coronaviridae epitope data was retrieved from the Immune Epitope Database and Analysis Resource (IEDB) on July 20th 2020^34^. The IBM Research Functional Genomics Platform (FGP)^35^ with updated SARS-CoV-2 genome annotation method^36^ was used to identify and retrieve the protein sequences, domain sequences and genome accessions from January 2020 to August 2020. Even though the data from early on in the pandemic, we show the relevance of results, specifically the stability of top 10 epitopes, for newer variant of concern, Delta, as well.

### 2.1 Immune Epitope Database

From IEDB, linear B-Cell and T-Cell epitopes for all Coronaviridae were downloaded. This data was filtered for *positive assays* only i.e. epitopes known through confirmed laboratory experiments. No filters for host, MHC restrictions, or diseases were applied. For epitope sequences in IEDB that were found originally in other Coronaviridae, the identification of those sequences in SARS-CoV-2 was based on observation of the exact sequence in SARS-CoV-2 proteins in FGP (see sections 3.2 and 3.3).

At the time of download, the data contained 601 linear T-cell epitopes with positive assays across 10 proteins and 23 organisms, and 586 linear B-cell epitopes with positive assays across 12 proteins and 30 organisms. There were 28 column descriptors of which the following were extracted for this study:

1. Epitope ID
2. Parent Protein
3. Parent Protein Accession
4. Antigen Name
5. Epitope Description (Sequence)

From IEDB, we retrieved all distinct (non-redundant) epitope sequences, the name of the protein from which they were found, and the antigen on which it was identified. This was used as ground truth data to locate conserved regions across SARS-CoV-2 proteins and to identify new epitopes not originally in IEDB.

Supplemental Figure S2 quantifies the data availability of IEDB B-Cell and T-Cell epitope by organism and protein. We don’t separate the epitopes based on MHC classes they bind to. This is because our methodology does not rely on prediction tools, where this separation can be important to select the correct allele and tools. Additionally, we wanted to see if clustering of epitopes happens based on MHC classes to which they belong. Also, MHC class can be easily deduced from the length of epitopes^37,38^.

### 2.2 SARS-CoV-2 Genomic Data

The Functional Genomics Platform (https://ibm.biz/functional-genomics) is a comprehensive database and analytics platform that provides (at the time of this work) ~300,000 pre-annotated bacterial and viral genomes, with over 75 million unique gene sequences, over 57 million protein sequences and over 263 million functional domains. For SARS-CoV-2, it contains high quality, curated genomic data defined as genomic sequences with <1% N’s (unknown bases), over 29000 bp and <0.05% unique amino acid mutations (as described in^36^) along with gene, protein, and functional domain sequences as well as KEGG, IPR and GO codes for those genomes. Moreover, it also provides analytical capabilities such as BLAST over the data. This data, as well as the analytical tools, can be accessed via the Python SDK or the API endpoints provided by FGP after creating a free account.

For this study, we utilized ~28K unique SARS-CoV-2 protein sequences and ~49K related protein domains and annotations identified from a collection of ~62K high quality SARS-CoV-2 genomes from NCBI GenBank^39^ and GISAID^40^ using the methods described in Beck, et al.^36^ (full protein and domain sequences can be found with that publication). The proteins and domains identified by FGP are related to the respective source’s genome accession. There are a total of 16 different proteins by name (including hypothetical protein) for SARS-CoV-2 with 11 proteins annotated, on average, per SARS-CoV-2 genome. FGP uses md5 hash of a sequence in order to construct its unique identifier (uid). Thus, each protein sequence has uid used to identify it.

The host metadata used in this work was downloaded from NCBI GenBank^39^ and GISAID^40^. Results for predicted epitopes for Replicase polyprotein 1a and Replicase polyprotein 1ab may not be accurate due to insufficient data.

### 2.3 Data Availability

All epitope data can be obtained directly from iedb.org. All protein sequence data can be obtained from either of the following sources:

- Functional genomics platform (ibm.biz/functional-genomics).
- Publication regarding FGP’s annotation of SARS-CoV-2 data^36^.

All genome metadata can be obtained from NCBI^39^ and GISAID^40^ directly.

For any additional data or for raw formatted files, please reach out to the corresponding author.

## 3 Methods

### 3.1 Protein Sequence Diversity Analysis

In the reference genomes for SARS-CoV-2, the copy number for each protein is 1^41, 42^. After de novo assembly and annotation of 61850 genomes we found evidence for copy number greater than 1 in only 584 genomes (< 1 %). Since some of the analyses described below depends on subsequent multi-sequence alignment to the reference, these genomes were omitted from the analysis.

### 3.2 Identification of Conserved Epitope Sequences

First, we were interested in finding known epitope sequences that are present in our protein sequence corpus. In this section, we utilize the set of 28K SARS-CoV-2 protein sequences from FGP to determine conserved epitope sequences from two subsets within IEDB data: a limited set of ‘ground truth’ SARS-CoV-2 epitopes, and epitopes observed in other Coronaviridae. To compare these sequences, the following steps were completed:

- Identification of whether an epitope occurs on a protein sequence and at what locus (amino acid position) using exact string matching functions provided in base Python. For this, the epitope’s parent protein was matched to the SARS-CoV-2 protein e.g. epitopeA, whose parent protein indicated in IEDB is Nucleoprotein, would be checked for presence on SARS-CoV-2 Nucleoprotein sequences only.
- Calculation of the number of times an epitope occurs on a protein sequence.

The above analysis was performed separately for B-Cell and T-Cell epitopes and for all protein names in the set of parent protein names retrieved from IEDB.

### 3.3 Identification of Candidate Epitope Sequences

Next, we wanted to detect candidate epitopes, i.e., potential peptides with high sequence homology to the laboratory confirmed epitopes. This would enable accounting for slight changes in known epitope sequences due to evolving protein regions.

To this end, we completed a sequence search using BLAST^43^ to compare all epitopes downloaded from IEDB against all corresponding (with same parent protein name) SARS-CoV-2 protein sequences using FGP’s BLAST service. FGP provides the capability to run BLAST against pre-constructed databases of nucleotide and amino acid sequences. The SARS-CoV-2 amino acid sequence database was selected for this study. Default parameters were used to run BLASTN and an e-value threshold of 0.01 was chosen since that indicated statistically significant match^44^.

We did not rely on generating candidate epitopes using epitope prediction tools since their accuracy with regards to SARS-CoV-2 may not be very high^37^.

The steps outlined in section 3.2 were repeated with the newly identified set of candidate epitopes to find all protein sequences where these epitopes were found, along with the start indices of those epitopes and their frequency per protein sequence. The results were appended to the output from conserved epitope sequence calculation and these were marked as CANDIDATE, and the conserved epitopes were marked as ORIGINAL.

### 3.4 Summary Statistics

Having compiled a list of ORIGINAL and CANDIDATE epitopes found in our set of 28k protein sequences, we wanted to study the presence of these epitopes on proteins and genomes. Specifically, we wanted to answer questions like: What are the most abundant epitopes? Is an epitope found multiple times in a protein sequence? How many epitopes are present on average in a genome?

### 3.5 Epitope Clustering and Classes

To explore how different epitopes relate to one another based on sequence homology and similarity of their loci on the protein sequence, we performed both sequence based clustering and position based clustering of epitopes.

For sequence-based clustering, the following steps were performed:

1. First, epitopes (both *ORIGINAL* and *CANDIDATE*) were grouped based on functional type (T or B) and parent protein.
2. For each group above, the Levenshtein edit distance measure was calculated (using Python) for every epitope-epitope pair. This yields a square matrix with axes labeled by epitope, and cell values as the distances between all pairs.
3. Linkage (Python scipy package) was then run on the edit distance matrix using ‘Euclidean’ metric and ‘Single’ method.
4. The resulting linkage matrix was then used to compute and plot a dendrogram.
5. We then define a cluster threshold where cluster members had a length normalized edit distance less than 1. We color any epitopes not clustering together in blue.

After clustering epitopes by sequence and extracting the relevant clusters for each protein and epitope type (B or T) combination, the occurrence of these clusters on SARS-CoV-2 genomes was investigated. To this end, the dendrogram was combined with a bar plot indicating frequency of occurrence of all epitopes in the cluster combined to which the epitope belongs.

A stacked bar plot was generated to present the distribution of genomes over three categories of original data source (hosts/samples): humans, animals, and environmental. Source data was retrieved from genome metadata files from NCBI and GISAID.

To obtain position-based clustering of the epitopes, the following steps were carried out:

1. B-Cell and T-Cell epitopes were divided into separate sets based on parent protein as described above.
2. To remove spurious sequences or assembly errors, we considered only sequences with length within ±10% of the UniProt45 reference protein sequence for SARS-CoV-2 e.g. for Nucleoprotein, the length of reference protein is 419 amino acids, and we allowed all sequences which were within a ± 10% range of that i.e. between 378 and 461 amino acids in length. For SARS-CoV-2 Spike glycoprotein, the reference protein is of length 1273 amino acids and allowed range was 1146 to 1400 amino acid characters.
3. Furthermore, we ran multi-sequence alignments (MSA) of all sequences per protein name relative to one another using MAFFT (v7.431) with the *-reorder* option^46^.
4. For each protein, a matrix was then constructed with the x-axis indicating the amino acid position on the protein and the y-axis indicating the epitopes where the length of the x-axis was the maximum length of the protein sequence.
5. Then each cell was assigned a binary value i.e. 0 or 1. The cell is assigned 1 if the epitope on that row is found on the position on protein corresponding with the column index of the cell. E.g. if epitopeA is on row0 and is found on a Nucleoprotein sequence between indices 7 and 15, then index 7 would be filled with 1.
6. After sequences have been aligned in the MSA, the positional information was padded so that all pairwise comparisons are represented with identical start and stop coordinates and gaps are filled in with dashes. To accommodate this relative positional information generated by an MSA, regular expressions were used to identify the start index of epitope sequences with the allowance for epitopes that span “gapped regions” due to the filling in of sequences in the MSA.
7. Epitope sequences spanning regions of insertion described in step 6 were investigated further for key non-synonymous mutations and their prevalence in this corpus of ORIGINAL and CANDIDATE epitopes.
8. After alignment the epitopes were found on the same position within the protein sequences. We colored the bars representing epitopes by their presence across the genomes.
9. After constructing the matrix in steps 2 and 3, ‘single’ linkage was run using method ‘Euclidean’ metric.
10. Finally, the clustermap and dendrogram were plotted using python’s seaborn library.

### 3.6 Identification of Epitopes on Protein

#### 3.6.1 Mutations in Proteins affecting Epitopes and Mutation Density

Certain epitopes were not found by MSA as exact subsequences on proteins. In such cases, we used a regular expression (regex) match to find the start position for the epitope. Additionally, these regions identify locations where protein mutations are occurring, and were thus studied in greater detail by visualizing MSA results.

Mutations at any given position on the spike glycoprotein transcript were logged using the MSA results as percentages. The proportion of amino acids matching the reference were measured at every position on the sequence to estimate “mutation density”. These were plotted to examine regions of elevated mutagenesis on the protein sequence with respect to the MSA-derived consensus sequence. Mutation density plots were constructed for both Spike glycoprotein and Nucleoprotein transcripts.

#### 3.6.2 Epitope Distribution Across Protein

The clustermap generated in 3.5 for position based clustering also serves to highlight where epitopes lie on the protein. However, to gain an amino acid-position level understanding of “immunodominance”, we also visualized the frequency of epitope occurrence at a given position on the protein transcript. Our epitope samples were split into ORIGINAL and CANDIDATE epitopes. We logged the number of times that an epitope occurred across all genomes. These frequencies were normalized and plotted along with the normalized global median frequency in order to assess regions of immunodominance relative to other regions on the protein transcript. The proportion of immunodominance prescribed to each position by new epitopes was drawn as a stacked area plot atop the proportion of immunodominance prescribed by original epitopes to compare the contributions of each epitope origin to the overall immunodominance level for the position. This graph was constructed for T-Cell and B-Cell epitopes on N and S proteins.

To identify epitope relationships within protein functional domains, the domains for each protein were also checked against epitopes for complete sequence homology i.e. if an epitope sequence exactly matches, or is found as a subsequence within a domain sequence for the corresponding protein.

#### 3.6.3 Epitope localization on protein structure

In order to visualize the position of the epitopes on the quaternary structure of the Nucleoprotein (N) and the Spike glycoprotein (S) of SARS-CoV-2 we used the VMD software47 and the following procedure for both proteins. First, we compared the reference sequence from UniProt with the sequence of the protein structures deposited in the Protein Data Bank^48, 49^ and then we visualize the epitope positions in the 3D shape of the protein by selecting the protein deposited structure showing 100% identity with the consensus sequence. Specifically, for the N protein we used the 1.5 Å resolution X-ray structure of the C-terminal domain (CTD) resolved by Y. Peng et al. ^3^ (PDB ID: 7CE0); for the S protein in the prefusion and postfusion conformations we used the cryo-EM structure resolved by Y. Cai et al.^21^ at 2.9 and 3.0 Å (PDB ID: 6XR8 and 6XRA respectively).

### 3.7 Result verification

We wanted to understand the correctness of our results and decided to use the latest set of lab confirmed epitopes to do the same. We did not rely on comparing results with other epitope prediction tools since their accuracy with regards to SARS-CoV-2 may not be very high^37^.

We analyzed the predicted set of epitopes against the latest set of epitopes from IEDB. We used BLAST to find the sequence alignment between the predicted and the known epitope sequences found on Spike glycoprotein and Nucleoprotein. For each protein, we first separated the epitopes found in B-Cell and T-Cell and created a BLAST database using the epitope sequences from IEDB. Thus, we generated four BLAST databases, one for each protein and cell type. We then performed a protein BLAST query with the predicted epitopes as query sequences. All default BLAST inputs were used, except the number of alignments was set to 1 in order to find the IEDB sequence that best matches each predicted sequence. For each protein and cell type, we then calculated the average percent identify and e-value between the predicted epitopes and known epitopes.

Additionally, even though a complete analysis with the latest variants of concern was out of the scope for this paper, in light of the outbreak and seriousness of Delta variant of SARS-CoV-2, we got 2 frequently seen sequences of Spike glycoprotein for Delta variant and checked if our top Spike T-Cell and Spike B-Cell epitopes are present on this Spike sequence. The sequences were obtained from36 and are included in Supplemental files SD1. As seen here^36^, these sequences have been found in other variants of concern as well.

## 4 Results

### 4.1 Protein Sequence Diversity Analysis

In this work, we analyzed a corpus of 61,850 SARS-CoV-2 genomes from NCBI GenBank and GISAID. Protein annotations for each were downloaded from FGP. After performing the data sanitation steps outlined in Methods Section 3.1, we identified 584 (< 1 %) genomes where after annotation, protein sequence diversity was >1 for one or more proteins. These were excluded resulting in a total genome dataset of 61,266. 4,595 and 1,737 unique protein sequences were observed in S and N sequences, respectively. From this corpus, the number of observed epitopes are indicated in Table 1.

**Table 1.** Epitope Distribution by Protein

| Protein Name | #Unique Sequences | B-Cell Epitopes |  | T-Cell Epitopes |  |
| --- | --- | --- | --- | --- | --- |
|  |  | Conserved | Candidate | Conserved | Candidate |
| Spike glycoprotein | 4595 | 50 | 152 | 114 | 29 |
| Nucleoprotein | 1737 | 52 | 62 | 65 | 54 |
| Membrane Protein | 285 | 8 | 16 | 18 | 9 |
| Protein 3a | 4 | 1 | 1 | 21 | 13 |
| Envelope Small Membrane Protein | 1 | 1 | 0 | 3 | 0 |
| ORF6 protein | 109 | 0 | 0 | 5 | 1 |
| Protein 7a | 217 | 0 | 1 | 9 | 3 |
| Protein 7b | 98 | 0 | 0 | 2 | 1 |
| Protein 9b | 7 | 0 | 1 | 0 | 0 |
| Replicase Polyprotein 1a* | 11509 | 0 | 4 | 3 | 0 |
| Replicase Polyprotein 1ab* | 795 | 0 | 0 | 39 | 0 |

### 4.2 Epitope Sequences

Section 3.2 outlines the procedure followed for identification of conserved epitopes between SARS-CoV-2 protein sequences and other Coronaviridae. As shown in Table 1, we identify 112 B-Cell and 279 T-Cell conserved epitopes. Of the 112 linear B-Cell epitopes from Coronaviridae found in SARS-CoV-2 protein sequences, 3 are originally from SARS-CoV-2 and 109 from SARS. Of the 279 linear T-Cell epitopes from Coronaviridae that were found in FGP SARS-CoV-2 protein sequences, 221 are originally from SARS-CoV-2 and 58 from SARS.

Additionally, we identified 492 candidate linear B-Cell(304) and T-Cell(188) epitopes in SARS-CoV-2 proteins with high sequence similarity to epitopes from IEDB(See Table 1). Our results suggest that linear SARS-CoV-2 epitope sequence length varies from ~7-42 amino acid characters for both ORIGINAL and CANDIDATE epitope sequences.

We also identify and list the top 10 B and T-Cell epitopes for Spike glycoprotein and Nucleoprotein in Tables 2 and 3. The criteria for being a top epitope is based only on the number of genomes an epitope is found in. In addition to the epitope sequences, the tables also list the start index of epitopes on aligned protein sequences, parent epitope (if the epitope is derived) and other epitopes which have high sequence similarity.

**Table 2.**
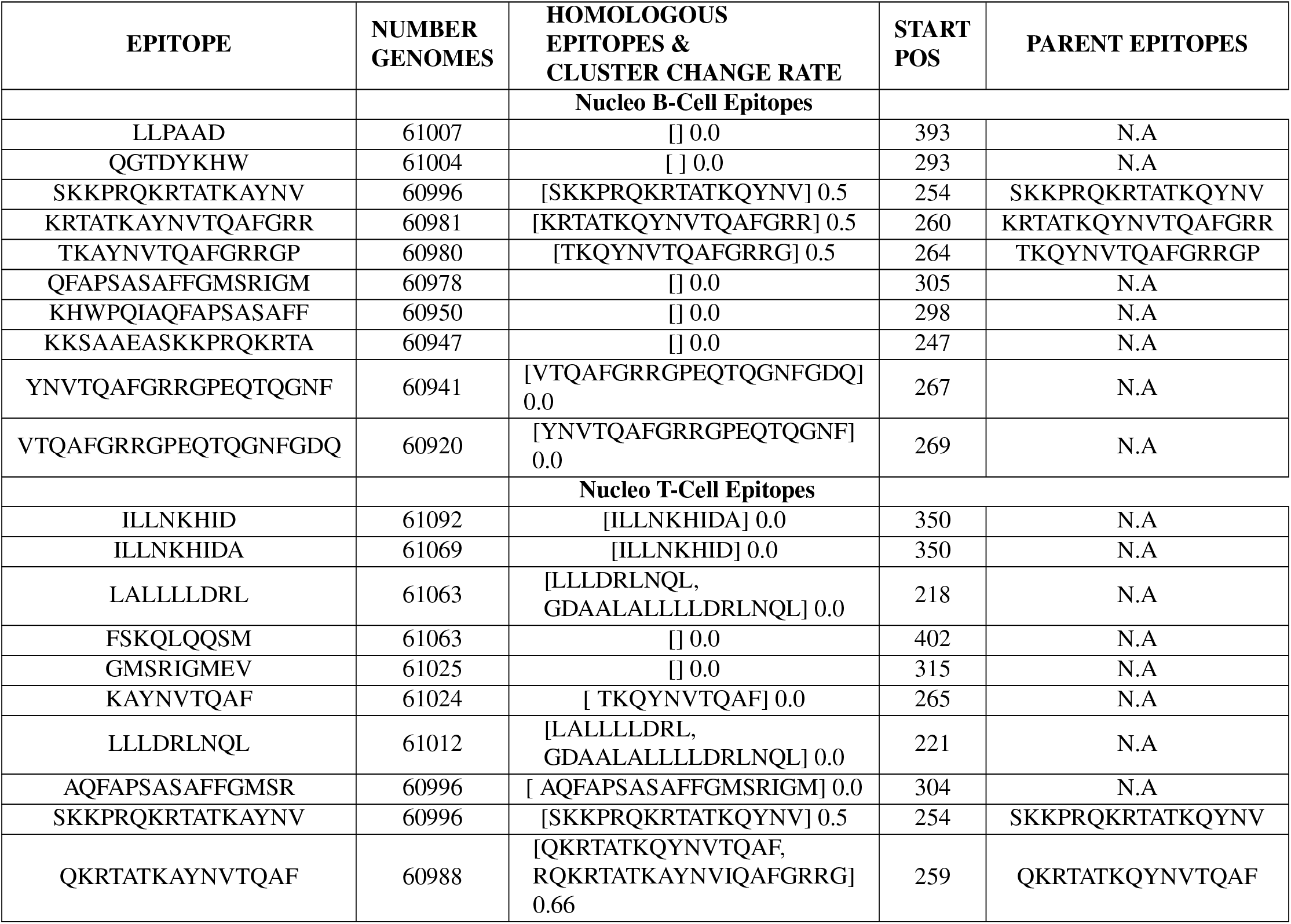
Top 10 most commonly found epitopes in Nucleoprotein. Here frequency of occurrence is determined by calculating the number of unique genomes an epitope is present in. Homologous epitopes are sister epitopes which clustering together with the epitope in first column when performing clustering by sequence. The cluster change rate indicates the probability of a candidate epitope being found in the cluster. If the cluster consists of only lab confirmed epitopes, then evolution rate will be 0. Start pos is the index of the starting position of epitope on aligned protein sequences as mentioned in 3.6.1 and in case of every epitope matches the median start position on unaligned sequences.

**Table 3.** Top 10 most commonly found epitopes in Spike glycoprotein sequences. Here frequency of occurrence is determined by calculating the number of unique genomes an epitope is present in. Homologous epitopes are sister epitopes which clustering together with the epitope in first column when performing clustering by sequence. The cluster change rate indicates the probability of a candidate epitope being found in the cluster. If the cluster consists of only lab confirmed epitopes, then evolution rate will be 0. Start pos is the index of the starting position of epitope on aligned protein sequences as mentioned in 3.6.1 and in case of every epitope matches the median start position on unaligned sequences.

| EPITOPE | NUMBER GENOMES | HOMOLOGOUS EPITOPES & CLUSTER CHANGE RATE | START POS | PARENT EPITOPES |
| --- | --- | --- | --- | --- |
| **Spike B-Cell Epitopes** |  |  |  |  |
| ILSRLDKVEAEVQIDRL | 61222 | [ILSRLDKVEAEVQIDRL]<br>0.0 | 979 | N.A |
| DFCGKGYHLMSFPQSAP | 61215 | [DFCGKGYHLMSFPQSAP]<br>1.0 | 1040 | DFCGKGYHLMSFPQAAP |
| MAYRFNGIGVTQNVLYE | 61213 | [MAYRFNGIGVTQNVLYE]<br>0.0 | 901 | N.A |
| VLGQSKRVDFCGKGYHL | 61212 | [VLGQSKRVDFCGKGYHL]<br>0.0 | 1032 | N.A |
| AISSVLNDILSRLDKVE | 61211 | [AISSVLNDILSRLDKVE]<br>0.0 | 971 | N.A |
| GSFCTQLN | 61209 | [GSFCTQLN]<br>0.0 | 756 | N.A |
| EAEVQIDRLITGRLQSL | 61205 | [EAEVQIDRLITGRLQSL]<br>0.0 | 987 | N.A |
| KQLSSNFGAISSVLNDI | 61205 | [KQLSSNFGAISSVLNDI]<br>0.0 | 963 | N.A |
| SLQTYVTQQLIRAAEIR | 61200 | [SLQTYVTQQLIRAAEIR]<br>0.0 | 1002 | N.A |
| LMSFPQSAPHGVVFLHV | 61199 | [LMSFPQSAPHGVVFLHV]<br>1.0 | 1048 | LMSFPQAAPHGVVFLHV |
| **Spike T-Cell Epitopes** |  |  |  |  |
| FPQSAPHGVVF | 61227 | [] 0.0 | 1051 | N.A |
| RVDFCGKGY | 61225 | [] 0.0 | 1038 | N.A |
| VLNDILSRL | 61225 | [SVLNDILSRL] 0.0 | 975 | N.A |
| SVLNDILSRL | 61224 | [VLNDILSRL] 0.0 | 974 | N.A |
| YHLMSFPQSA | 61223 | [] 0.0 | 1046 | N.A |
| MAYRFNGIGVTQNVLY | 61216 | [] 0.0 | 901 | N.A |
| ALNTLVKQL | 61212 | [AQALNTLVKQL] 0.0 | 957 | N.A |
| AQALNTLVKQL | 61211 | [ALNTLVKQL] 0.0 | 955 | N.A |
| SFPQSAPHGVVFLHV | 61210 | [LMSFPQSAPHGVVFLHV] 0.5 | 1050 | N.A |
| LITGRLQSL | 61209 | [] 0.0 | 995 | N.A |

In order to test our predicted epitopes, and to measure the sequence diversity, we compared the predicted epitope sequences to the known epitopes for each cell type and calculated the percent match that is found. We find the average percent identity for Nucleoprotein B-Cell is 95.05%, Nucleoprotein T-Cell is 97.75%, Spike B-Cell is 94.64%, and Spike T-Cell is 97.40%. We also analyzed the range of percent identities for all predicted epitopes within each cell type. Figure S1 shows the average and range of percent identity for each epitope class. This shows that while the average for all 4 classes are greater than 90%, the sequence diversity of B-Cell epitopes is larger than for T-Cell epitopes.

Additionally, to verify the validity of the top Spike epitopes in the Delta variant, we checked the presence of top 10 B and T-Cell Spike epitopes on 2 SARS-CoV-2 Delta spike sequences. We found that all epitopes are indeed present on the Delta Spike as well.

Supplemental Figures S19, S20, S21, S18, S16, S17 show presence of epitopes in genomes and proteins. We observe an almost bi-modal distribution. This may require further investigation into genomes with low number of identified epitopes, since these genomes may have sequencing errors or other assembly defects etc. Note that each epitope was found only once on a protein sequence, and each protein had a copy number of 1 on a genome, thus each epitope was found at most once per genome (if found at all).

Figures 1, S5, S3, S4, show the relations between epitopes, from sequence based clustering.

**Figure 1.**
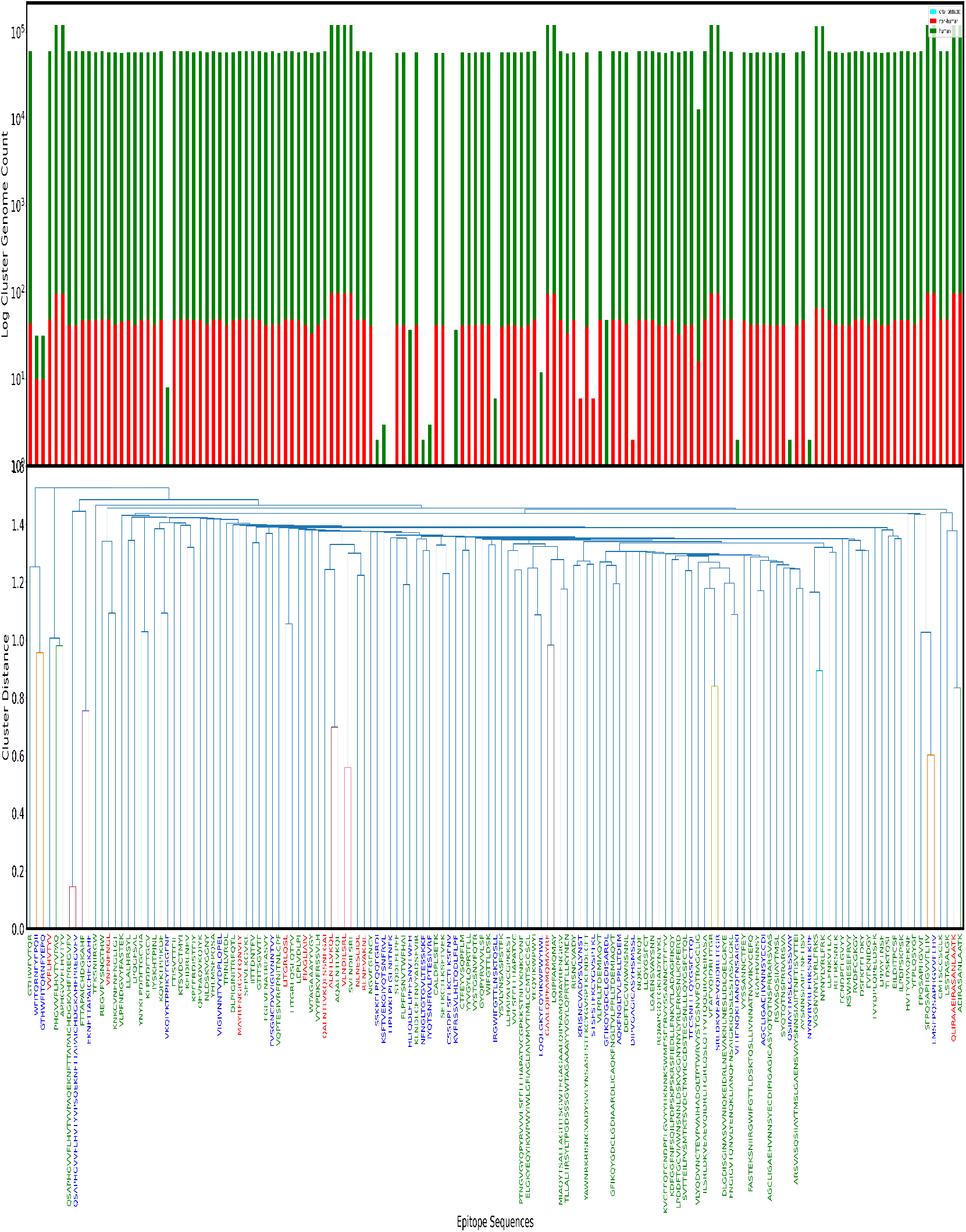
Clustering of Spike glycoprotein T-Cell Epitopes *ORIGINAL* + *CANDIDATE* and their occurrence. The bottom chart is a dendrogram obtained by performing sequence based clustering on T-Cell S epitopes. The labels along x-axis in dendrogram are the epitope sequences and have been assigned colors based on their originating organism, Blue if CANDIDATE epitope, RED if originally found in SARS and GREEN if known to be found in SARS-CoV-2. Along the y-axis of the dendrogram is the edit distance score. Edit distance of two sequences lets us know how similar the sequences are to one-another. We put a threshold of 1.0 on this edit distance to discover clusters within the epitopes i.e. epitopes with normalized distance <1.0 are part of same cluster. In top part of each figure, the bars align with the epitope labels from dendrogram. Each bar represents the number of times all members of the cluster, to which the epitope belongs, are seen across SARS-CoV-2 genomes in our dataset. It is also important to note that the figure is actually a log-log plot of the counts. Furthermore, each bar is stacked based to show genomes sequenced in humans, animals or environment. We would also like to highlight that low presence in genomes sequenced from environment is not a consequence of epitopes not being found in those genomes, but rather a product of extremely low number of high quality genomes from environment in our dataset.

We observe that in Fig 1 most epitopes are already laboratory confirmed epitopes in SARS-CoV-2. We observe only one cluster with significant frequency which has more than two epitopes. In Supplemental Fig S5 for Spike B-Cell, we see most epitopes are CANDIDATE epitopes indicating high sequence diversity and potentially many undiscovered or unconfirmed epitopes at the time of analysis. There is only one cluster with significant presence, and even within that there is only one laboratory confirmed epitope. In Supplemental Fig S4 for Nucleoprotein B-Cell epitopes, we observe that there are no laboratory confirmed epitopes already known to be present in SARS-CoV-2 and top clusters have non-zero CANDIDATE epitopes. This may indicate rapidly evolving regions on proteins leading to evolution of new epitope sequences. In Supplemental Fig S3 for Nucleoprotein T-Cell epitopes, we see that the cluster with the most significant presence has only SARS and CANDIDATE epitopes.

In all figures above, we observe a significant number of CANDIDATE epitopes. When we compare the presence of clusters against each other, we observe that there are certain clusters of epitopes which have a much higher weight than other clusters.

### 4.3 Epitopes on Protein

#### 4.3.1 Mutations in Proteins affecting Epitopes

Understanding the rate of mutation across epitope sequences can provide insights into waning host immunity, and the average period of host reinfection. These insights can also shape our understanding of vaccine efficacy over time. To evaluate this, we computed a multiple sequence alignment(MSA) of unique Spike glycoprotein and Nucleoprotein sequences and evaluated the mutational density across the B-Cell and T-Cell epitopes.

In Spike glycoprotein, Fig 3, we observe specific residue positions which have high mutation density when evaluating the consensus sequence obtained after running MSA. We also observe that a small region between residue positions 250-350 has slightly higher mutation density on average. In Nucleoprotein, we see slightly above average mutation density about residue position 200.

For Nucleoprotein, there were no epitopes that overlapped with mutations or indels (insertions or deletions). However for the Spike glycoprotein consensus sequence logo, the most prevalent amino acid by position was observed to be present with a median value of 99.84% (range 0.023%–100%) suggesting high sequence conservation across the protein. However, there was a small amount of mutations or indels in epitope sequences. For example, for epitope WTAGAAAYYVGY at amino acid position 272–288, we observed several key differences from the wild type epitope sequence. We observed a three amino acid insertion in one of our Spike glycoprotein sequences (occurring in X% of sequences). Additionally, at amino acid position 278, we observe higher variability where the dominant amino acid glycine (G) is present in only 91.41% of sequences with substitutions to serine (S) or aspartic acid (D) being the most common. This shifts from an aliphatic amino acid to an polar, hydroxylated amino acid or negatively charged acid, respectively, which can then change binding affinity to host proteins.

#### 4.3.2 Epitope Distribution Across Protein

The immunodominance Figures 3a, S13, S9, S12 show the presence of T-Cell and B-Cell epitopes on Spike glycoprotein and Nucleoprotein proteins. As discussed in section 3.6.1, we plot the epitopes on each protein and cluster them by position. Figure 2 shows the position of T-Cell epitopes on S. As we move top down, the median position gradually shifts from the C-terminus to the N-terminus of the protein. These plots relay more granular information regarding each epitope and it’s presence in our genome set and on the protein, along with it’s positional neighbors, as compared to immunodominance plots which show aggregate data for all epitopes. Where the slope is high the epitopes are more diverse and changing most rapidly.

**Figure 2.**
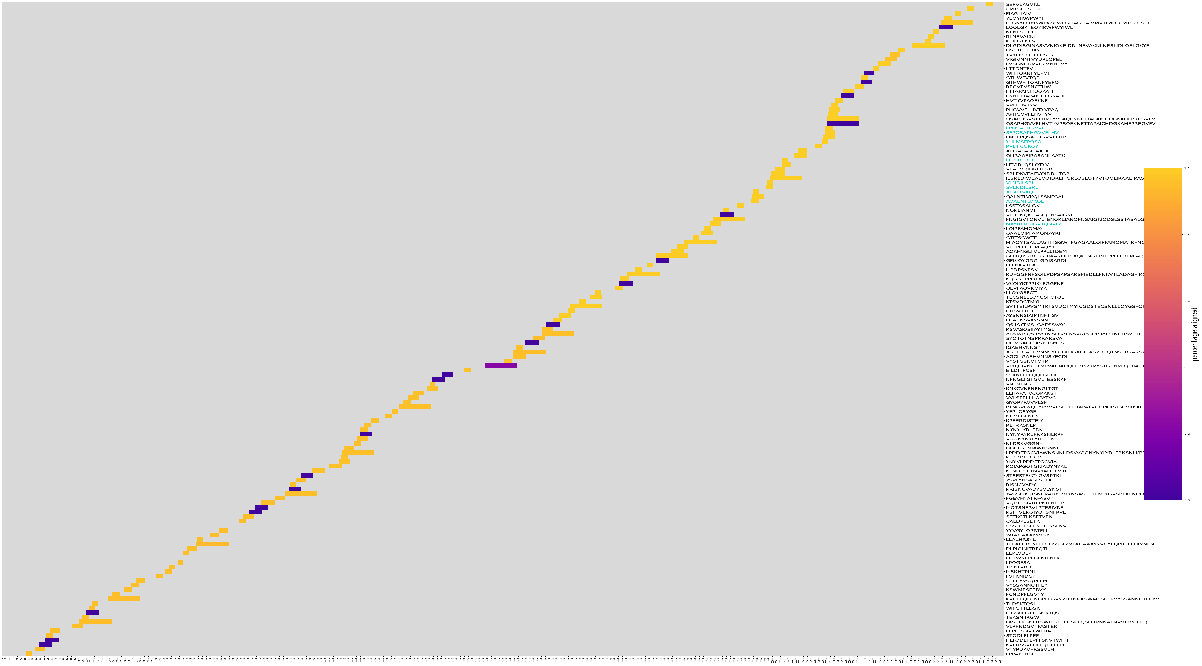
Clustermap obtained after clustering the T-Cell epitopes of Spike glycoprotein based on the position at which they occur within the protein. X-axis is the entire length of the protein, which is 1299 in case of S. Along the y-axis, every row represents one epitope. The color scheme is defined by using a color map which assigns colors to each row depending on occurrences of the epitope across all genomes. Y-axis labels on right hand side are colored cyan to indicate an epitope from top 10 list.

**Figure 3.**
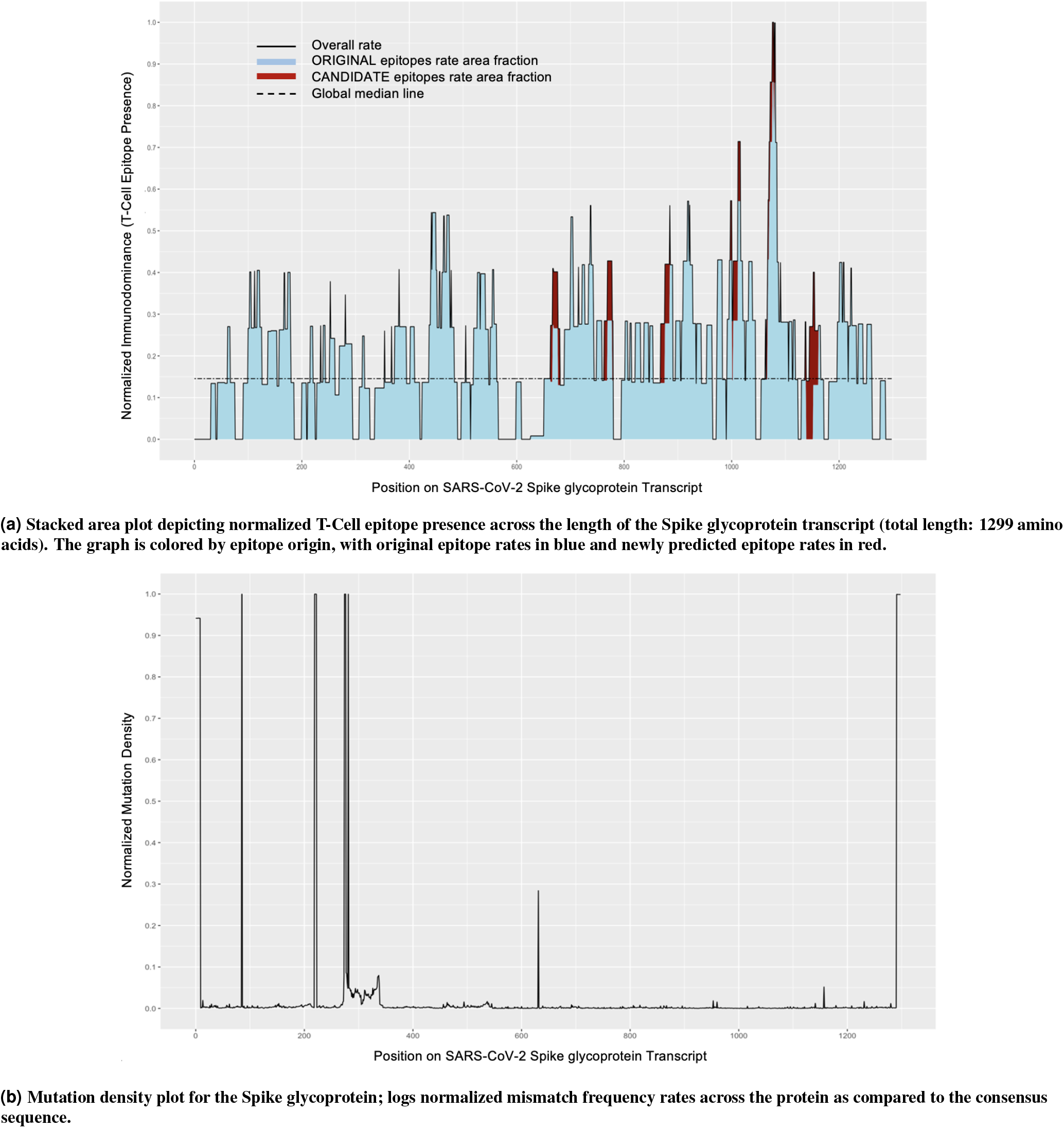
Immunodominance plot and mutagenesis plots.

In Fig 3a, we observe an almost even distribution of T-Cell epitopes on Spike glycoprotein, however we observe an uptick at 1063. We also notice fewer CANDIDATE epitopes as already noted in 4.1. By simultaneously looking at Fig 2, we observe that most prevalent T-Cell epitopes on Spike are in S2, and 3 contribute to the uptick along 1063.

Supplementary Fig S9 reveals a large number of CANDIDATE B-Cell epitope sequences on Spike glycoprotein and also significant regions on S1 with no epitopes present. This is also observed in Supplementary Fig S8, however is not as clear. From the latter we can again see that most prevalent Spike B-Cell epitopes are found on S2 region.

For Spike, presence of both T-Cell and B-Cell epitopes along S2 correlates with lower mutation density in S2.

Supplementary Fig S12 shows Immunodominance for Nucleoprotein T-Cell. The figure reveals that most ORIGINAL epitopes lie between residue positions 300 and 365 which is the C terminal (CTD). However, a considerable number of CANDIDATE epitopes are observed between residue positions 50 and 120 which is the N terminal domain. We also see few regions of gaps with fewer epitopes around residue position 200 which has high mutation density. From Supplementary Fig S6 we see that most prevalent epitopes are on the C terminal domain.

Immunodominance figure of Nucleoprotein B-Cell, Supplementary Fig S13 reveals that many CANDIDATE epitopes are found, even in regions where no ORIGINAL epitopes are present. We also observe a gap around residue position 200 which is the linker region. Most prevalent epitopes are again present in the C terminal domain, see Supplementary Fig S7.

To better understand the relationship between epitopes and protein functional domains, we evaluated the occurrence of complete sequence identity between epitopes and protein domains. Although epitopes are observed to span the length of Nucleoprotein and Spike glycoprotein, we did not observe any epitopes that exactly match a full length domain sequence or subsequence of any domain sequence on SARS-CoV-2 proteins. All matches found contained at minimum a single amino acid mutation. Domains and epitopes were only compared with respect to the same parent protein.

#### 4.3.3 Epitope localization on protein structure

We analyzed the position of the 10 B and T-Cell epitopes that were most frequently found in all genomes investigated, and we noticed that almost all of these high-frequency epitopes were localized in the C-terminal domain (C-terminal domain) of the N protein. Therefore, in Figure 4B-C we only show the C-terminal domain 3D structure of the N protein and indicate in Figure 4 A the position of the four epitopes located in the linker or in the C-terminal domain intrinsically disordered region (IDR) of the N protein. For greater clarity, we represent the position of B and T-Cell epitopes only on one homodimer, and have grayed out the others. The Nucleoprotein C-terminal domain structure consists of one 3_10_ helix followed by four *α*-helices, two *β*-strands, another *α*-helix and another 3_10_ helix^3^. In B-Cells, the 10 most frequently identified epitopes are located between the first 3_10_ helix and the first *β*-strand as shown in Figure 4B. The KKSAAEASKKPRQKRTA (bright blue) and the SKKPRQKRTATKAYNV (green) epitopes are on the first 3_10_ helix and overlap. The KRTATKAYNVTQAFGRR (red) epitope is on the first *α*-helix and is followed by the VTQAFGRRGPEQTQGNFGDQ (pink) one. Their sequence includes two other epitopes: the TKAYNVTQAFGRRGP (purple) and the YNVTQAFGRRGPEQTQGNF (cyan). The QGTDYKHW (orange) is located between the second and third *α*-helix followed by the KHWPQIAQFAPSASAFF (dark blue) and the QFAPSASAFFGMSRIGM (violet) epitopes, which extends half way of the first *β*-strand. Only one epitope, LLPAAD (yellow) among those represented here is located in the C-terminal IDR (Figure 4A,B). Compared to our observations for the B-Cell epitopes, the T-Cells the epitopes are distributed in a discontinuous manner in the C-terminal domain sequence. In addition, two epitopes are located in the linker (LALLLLDRL (green) and LLLDRLNQL (dark blue) respectively) and one in the C-term IDR (FSKQLQQSM (red)) (Figure 4A,C). The SKKPRQKRTATKAYNV (cyan) epitope is on the 310 helix and is followed by the KAYNVTQAF (violet) epitope located on the first *α*-helix. The sequence of these two epitopes includes that of the QKRTATKAYNVTQAF (pink). The AQFAPSASAFFGMSR epitope (bright blue) is on the third *α*-helix and partially overlaps the GMSRIGMEV (purple) epitope, which extends over almost all the first *β*-strand. The two ILLNKHID and ILLNKHIDA epitopes (yellow and orange) located on the last *α*-helix overlaps except for one residue.

**Figure 4.**
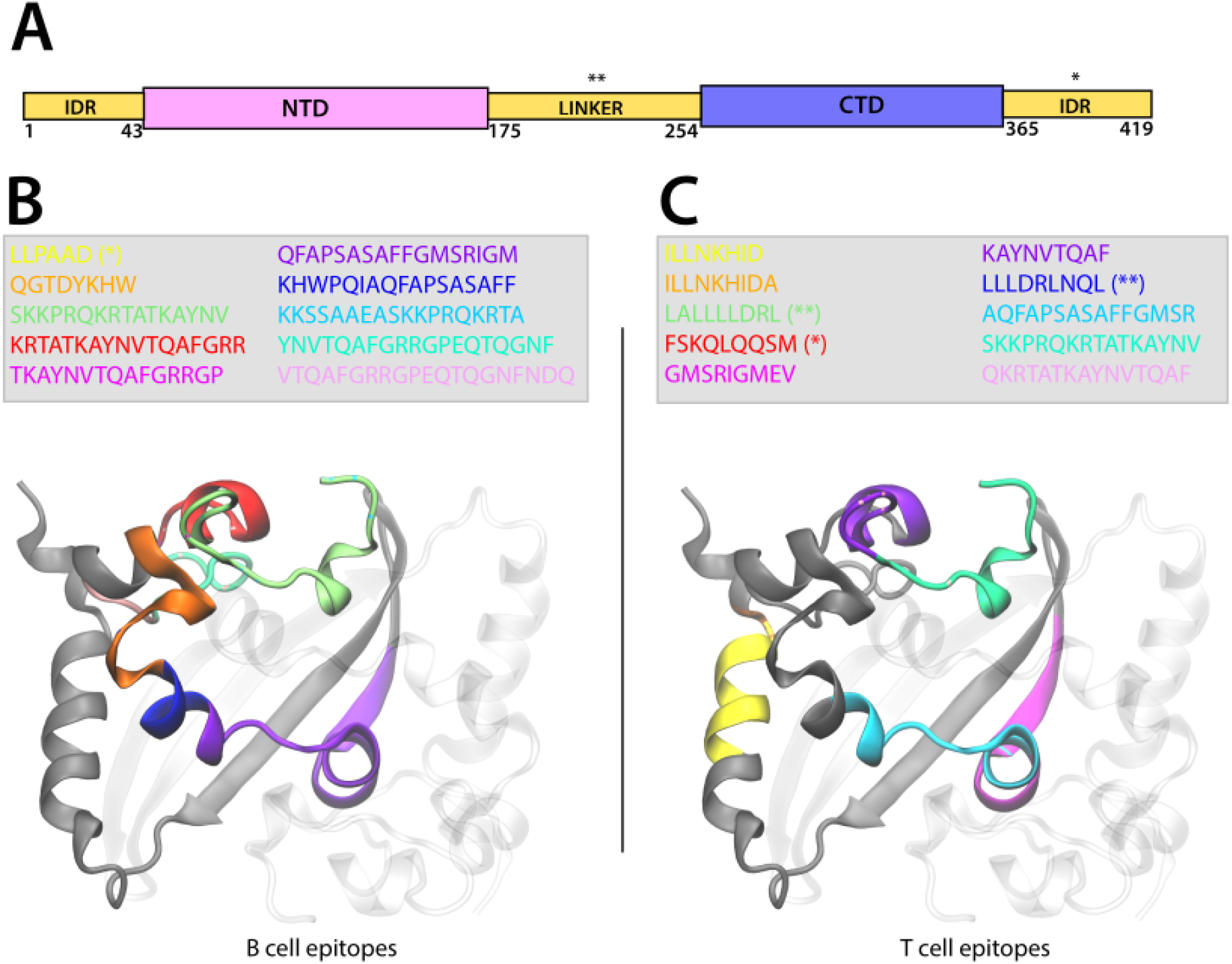
Representation of the localization of the B-Cell and T-Cell epitopes on the CTD domain of the Nucleoprotein. **A**. Scheme of SARS-CoV-2 N domains illustrating the N-term intrinsically disorder region (IDR) followed by the N-terminal domain (NTD), the IDR linker, the C-terminal domain (CTD), and the C-term IDR. **B-C**. The N CTD dimer is represented in New Cartoon format (one monomer is gray colored and the other is transparent) and the sequence of the B-Cell (**B**) and T-Cell (**C**) epitopes is colored according to the legend represented in the figure. The epitope sequence is represented in the legend. The epitopes located in the linker domain are indicated by (**) and those in the C-term IDR by (*). For great clarity, we represented the epitopes in only one monomer.

Because the S protein undergoes major conformational changes allowing membrane fusion between the SARS-CoV-2 viral membrane and the host cell, we show in the 3D representation of the S protein the position of the ten most frequently found B-Cell and T-Cell epitopes in all genomes investigated for both the prefusion and postfusion states Fig 5B-C. For clarity, we represented the epitopes only on one homotrimer and grayed out the rest. In addition, in Fig. 5A we show a schematic representation of the S protein sequence to identify visually the position of the epitopes on the protein.

**Figure 5.**
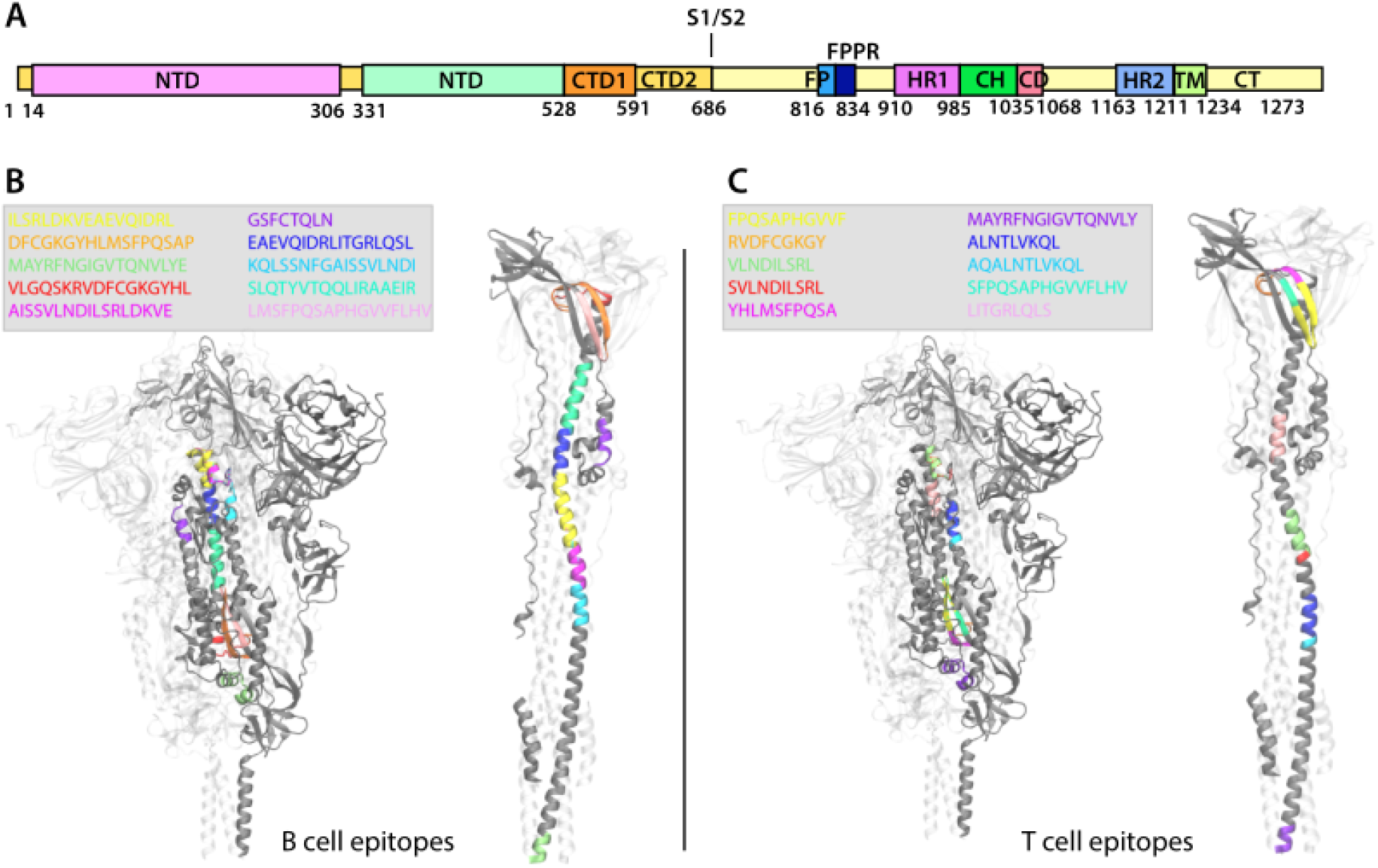
Representation of the localization of the B-Cell and T-Cell epitopes on the SARS-CoV-2 Spike glycoprotein in the prefusion and postfusion conformations. **A**. Scheme of SARS-CoV-2 S1 and S2 units of the S protein and of their domains. **B-C**.The S protein trimer is represented in New Cartoon format (one monomer is gray colored the other two are transparent) and is shown in the prefusion conformation in the left side of the panels and in the postfusion conformation on the right side of the panels. The sequence of the B-Cell (**A**) and T-Cell (**B**) epitopes is shown in the figure legend and is colored accordingly in the S protein structure.

We observe that the 10 B and T-Cell epitopes are localized in the S2 subunit of the S protein and specifically they are found between residue 902 and 1065 except the B-Cell epitope GSFCTQLN (violet) located between residue 757 and 764 before the fusion peptide (FP). The B-Cell epitope MAYRFNGIGVTQNLVYE (green) partially located in the heptad repeat 1 (HR1) is also recognized by T cells although the two epitope sequences differ for the glutamic acid E, which is absent in the T-Cell epitope sequence (violet colored sequence in Fig 5C). The five consecutive B-Cell epitopes KQLSSNFGAISSVLNDI (bright blue), AISSVLNDILSRLDKVE (purple) ILSRLDKVEAEVQIDRL(yellow), EAEVQIDRLITGRLQSL (dark blue), and SLQTYVTQQLIRAAEIR (cyan) are localized between the HR1 domain and almost all the central helix (CH) spanning between residue 964 and 1019 of the S protein. The remaining three B-Cell epitopes VLGQSKRVDFCGKGYHL (red), DFCGKGYHLMSFPQSAP (orange), and LMSFPQSAPHGVVFLHV (pink) are mostly localized in the connector domain (CD).

As we observed in the case of the N protein, the T-Cell epitopes are more widely distributed over the entire length of the S protein S2 subunit. The two epitopes ALNTLVKQL (dark blue) and VLNDILSRL (green) are located in the HR1 domain and partially overlap with the other two epitopes AQALNTLVKQL (bright blue) and SVLNDILSRL (red). The epitope LITGRLQSL (pink) is localized entirely in the CH domain. Consecutively located on the CD domain are the following epitopes: RVDFCGKGY (orange), YHLMSFPQSA (purple), FPQSAPHGVVF (yellow), and SFPQSAPHGVVFLHV (cyan).

## 5 Discussion

The ground truth epitope data contains many more entries from certain parent organisms, namely SARS and SARS-COV-2, and most often for the spike glycoprotein. Even though this is just a reflection of what was sequenced, it is important to remember this data bias whilst considering the results described in this paper, since it may influence the observed similarity with respect to other proteins in the SARS-COV-2 genomes.

Also, observe, that complete sequence homology is found only with respect to identified SARS-COV-2 and SARS epitopes. No epitopes from other Coronaviridae were found to be present in SARS-COV-2 genomes when performing an *exact sequence match*. This negative result is quite interesting and underscores the similarity between SARS and SARS-COV-2 genomes, and lack of similarity to other Coronaviridae, as has been observed in other studies of the COVID-19 virus^1^.

However, even though no exact match of epitopes was found with Coronaviridae other than SARS, when considering fuzzy matches or CANDIDATE epitopes, we see that 14 B-CELL epitopes had parent epitopes found in organisms other than SARS and SARS-COV-2. These organisms are: Murine hepatitis virus strain JHM, Feline infectious peritonitis virus (strain KU-2), Infectious bronchitis virus, Porcine epidemic diarrhea virus, Murine hepatitis virus strain A59,Avian infectious bronchitis virus (strain M41), Avian infectious bronchitis virus (strain Vic S), Porcine transmissible gastroenteritis coronavirus strain Purdue. Additionally, 26 NEW T-CELL epitopes were also found to have parent epitopes belonging to organism other than SARS and SARS-COV-2. These organisms are: Feline infectious peritonitis virus (strain KU-2), Human betacoronavirus 2c EMC/2012, Feline infectious peritonitis virus (strain 79-1146), Murine hepatitis virus, Avian infectious bronchitis virus (strain Vic S). This similarity between regions of SARS-CoV-2 proteins and immune targets of other Coronaviridae might also be interesting in scenarios concerning looking for viable therapeutics, refining epidemiological models to better estimate effective rate of transmission by accounting for possibly resistant population etc. The results shown in Supplementary Fig S1 provide validation for these epitope predictions. Also, since top 10 epitopes are found on SARS-CoV-2 Delta variant’s Spike glycoprotein sequence, this increases our confidence further in our predictions.

Tables 2 and 3 provide a list of the most commonly observed epitopes. For brevity we only discuss the top 10 epitopes for each epitope type and protein. Amongst the most frequent epitopes, we see that all derived epitopes have parent epitopes belonging to SARS except for one T-Cell Spike glycoprotein epitope, ISSVLNDILSRLDKVEAEVQ, which has a parent epitope from Feline infectious peritonitis virus *strain*79 – 114. The bulk of the most commonly seen epitopes are actually ORIGINAL epitopes i.e. lab confirmed epitopes found as is in SARS or SARS-COV-2 genomes.

Looking at epitope distributions in genomes from Supplemental Figures S18, S19 and number of epitopes present in a genome from Supplemental Figures S16, S21 we observe a stability of epitope presence across a large set of genomes which is a positive signal for vaccine development and effectiveness.

From our methods and results, four approaches emerge for selection of epitopes for vaccine design:

1. From sequence based clustering plots like Fig 1, we can also measure an evolution rate for a cluster by looking at the ratio of CANDIDATE epitopes to total epitopes in a cluster. This metric can be useful when evaluating epitope candidates for vaccines and can be used to theoretically predict the probability of change of an epitope solely on sequence homology. Additionally, mapping cluster presence across genome set adds a dimension to identifying the most suitable epitopes. We could also filter by host to study any changes that might arise from the virus evolving in different hosts. Tracking of major clusters could also enable development of statistical models to estimate a timeline for vaccine robustness.
2. Analyzing mutation density regions and immunodominance regions in order to evaluate which segments of proteins may be undergoing the fewest amino acid changes and thus would advise the most viable vaccine targets. This combined with studying position based clustering plots could add more insight for selection of epitopes since we could highlight most prevalent epitopes and also take into account their neighbors.
3. Figures 4 and 5 illustrate the position of the ten most commonly observed epitopes on the 3D representation of the N and the S proteins. We found that in the N protein, the epitopes are mostly localized in the CTD and span different stretches of amino acids depending on the type of cell. In the S protein, the epitopes are found essentially in the S2 subunit, which overall shares 91% amino acid sequence identity with the SARS-CoV S1 subunit. We found that the epitopes are primarily found in the HR1, CH, and CD motifs except for one B-cell epitope located in the region upstream the FP. Compared to prefusion conformation of the S protein, the S postfusion conformation seems to favor a greater exposure of the epitopes to the solvent and thus to antibody binding. The S protein is characterized by a highly density glycan surface, which can lead to immune evasion already studied in SARS-CoV-2 and other coronaviruses^50-53^. X. Fan et al.54 identified the putative sites of the N-linked glycans shielding the postfusion S protein surface. These sites are completely conserved between SARS-CoV-2 and SARS-CoV. The combined knowledge of the position of the most commonly observed epitopes and the glycan sites is crucial for developing broad-spectrum vaccines and therapeutics being the S protein the major determinant for viral transmission.
4. Stability of epitopes across different genomes and strain when data is available for the same.

There are important syntergies between to work of Grifoni et al.^1^ and the investigation reported here. They performed their work in early 2020 and discuss prediction of immune targets *in the absence of lab confirmed targets* to speed up vaccine design. Their approach is based on identifying protein and genome similarities to other notable Coronaviridae which have led to previous epidemics and pandemics including SARS-CoV-1 and MERS. Our work confirms some of the results presented by Griffoni et al. In particular both studies highlight the similarity between SARS-CoV-1 and SARS-CoV-2. Furthermore, some of the epitopes predicted by Griffoni et al have also been predicted and observed found in the current work. We would also like to highlight a few key differences: 1. The current analysis covers a much larger set of genomes. 2. At the time of our analysis, there were lab confirmed epitopes for SARS-CoV-2. 3. Instead of relying only on epitope prediction tools, we rely on exact matching with high sequence identity based approaches. 4. Griffoni et al. map epitopes on SARS-CoV-1 and study sequence identity between different regions of SARS-CoV-1 and SARS-CoV-2 to ultimately uncover regions with high epitope presence. In contrast this work relies directly on lab confirmed epitopes only (instead of mapping immunodominant regions).

## 6 Future Work

1. We could study the proximity of epitopes to more mutation prone regions to eliminate immune targets which otherwise may seem promising.
2. We could also combine the epitope evaluation approaches discussed in Section 5 to design a quantitative metric for evaluating and ranking epitope targets suitable for vaccine development. Selection of stable epitopes is important since that would enable development of shorter vaccines and may have a positive impact on effectiveness of vaccines since more stable immunotargets may be learned by the host immune system.
3. Studying the presence of top 10 epitopes in vaccine sequences.

## 9 Author Contributions

AA conceived of this work. AA and JK designed the experiments. AA, KLB, GN, GM, SC, JK and MK generated and analyzed the data. AA, KLB, GN, GM, SC, JK and SB wrote the manuscript. ES is the architect of the platform used. MK and VM provided scientific guidance and domain knowledge.

## Supplement

**Figure S1.**
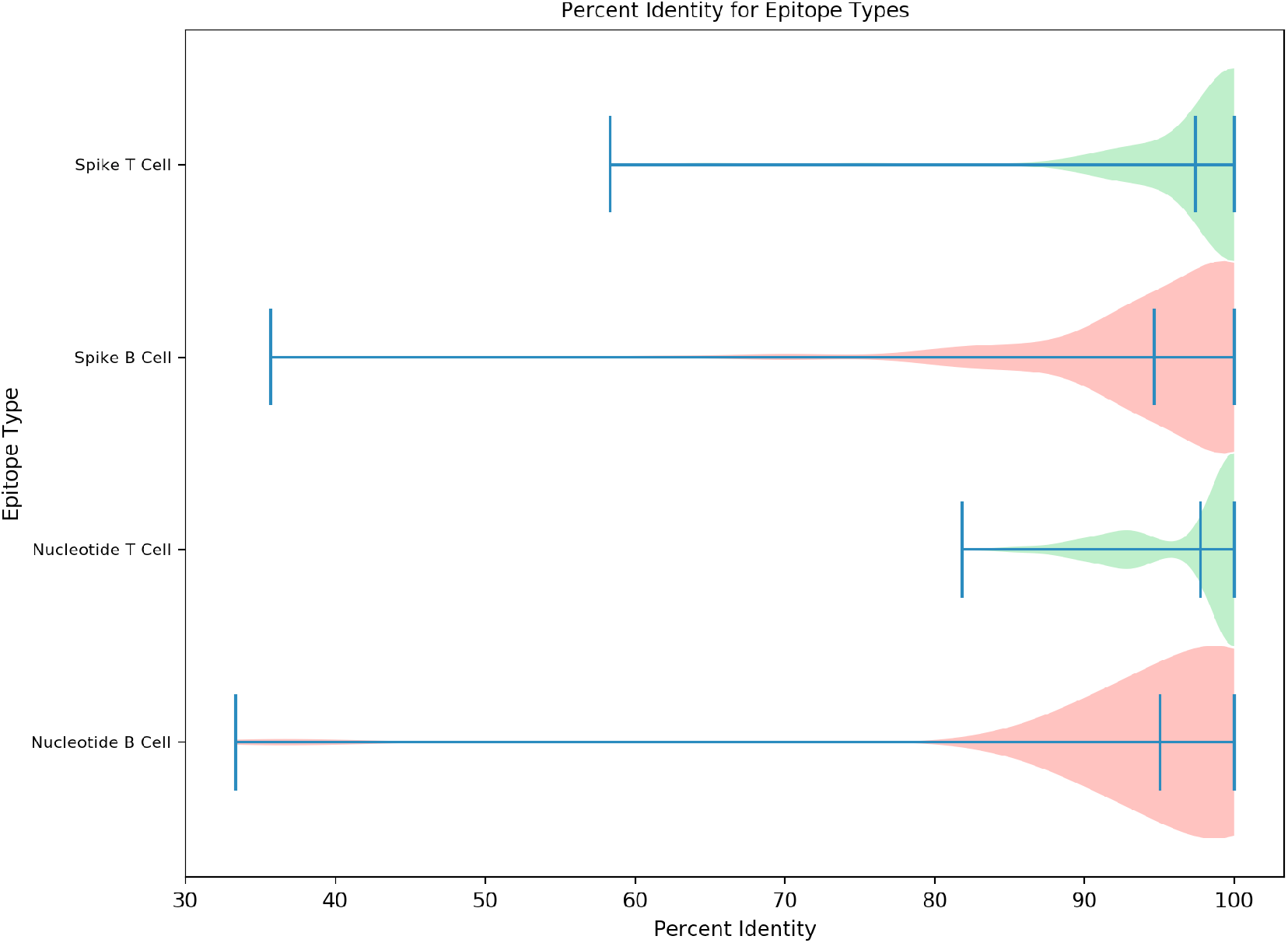
Percent identity of matches between predicted and known epitopes for each cell class.

**Figure S2.**
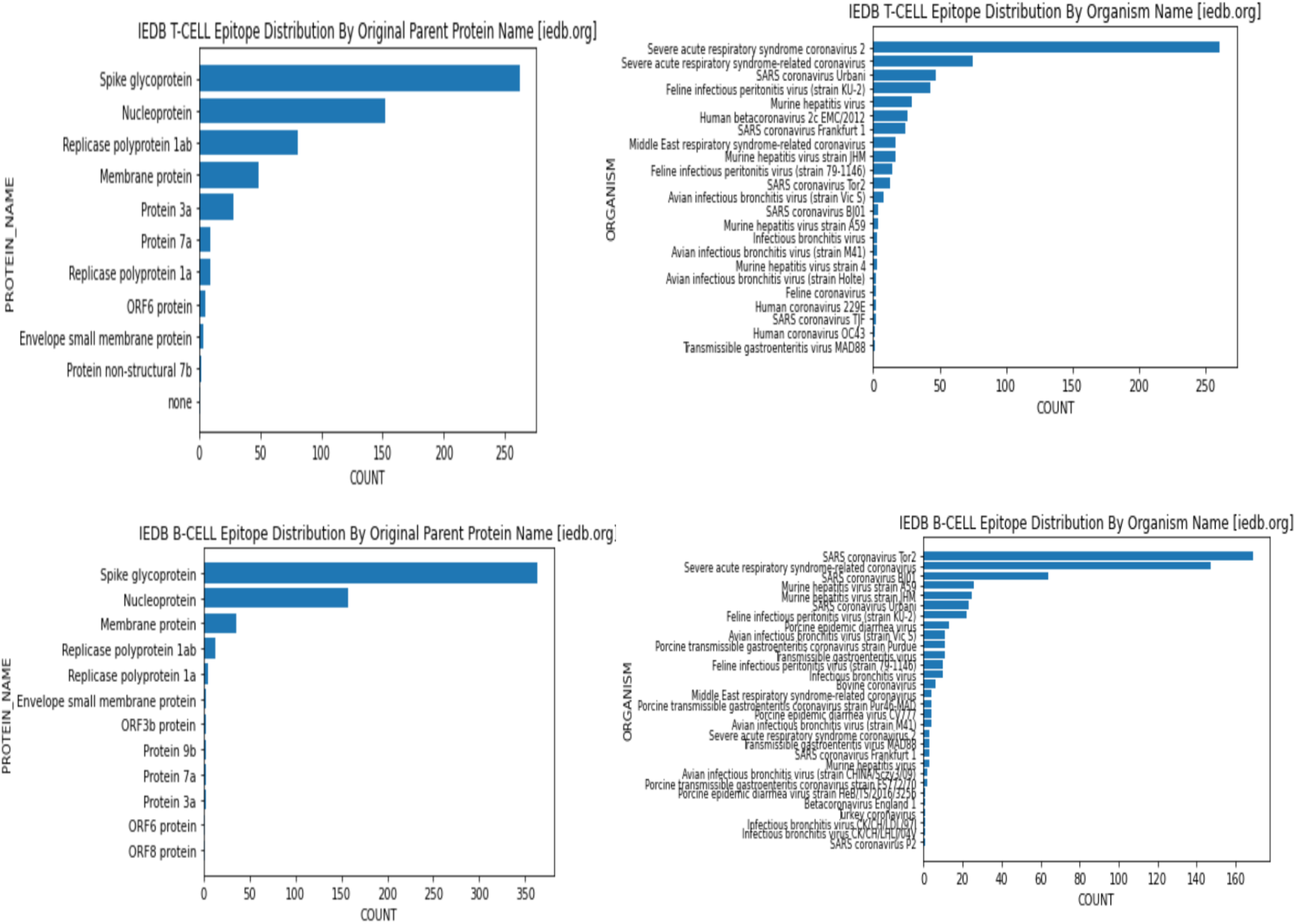
Distribution of Epitopes by Organism and Protein

**Figure S3.**
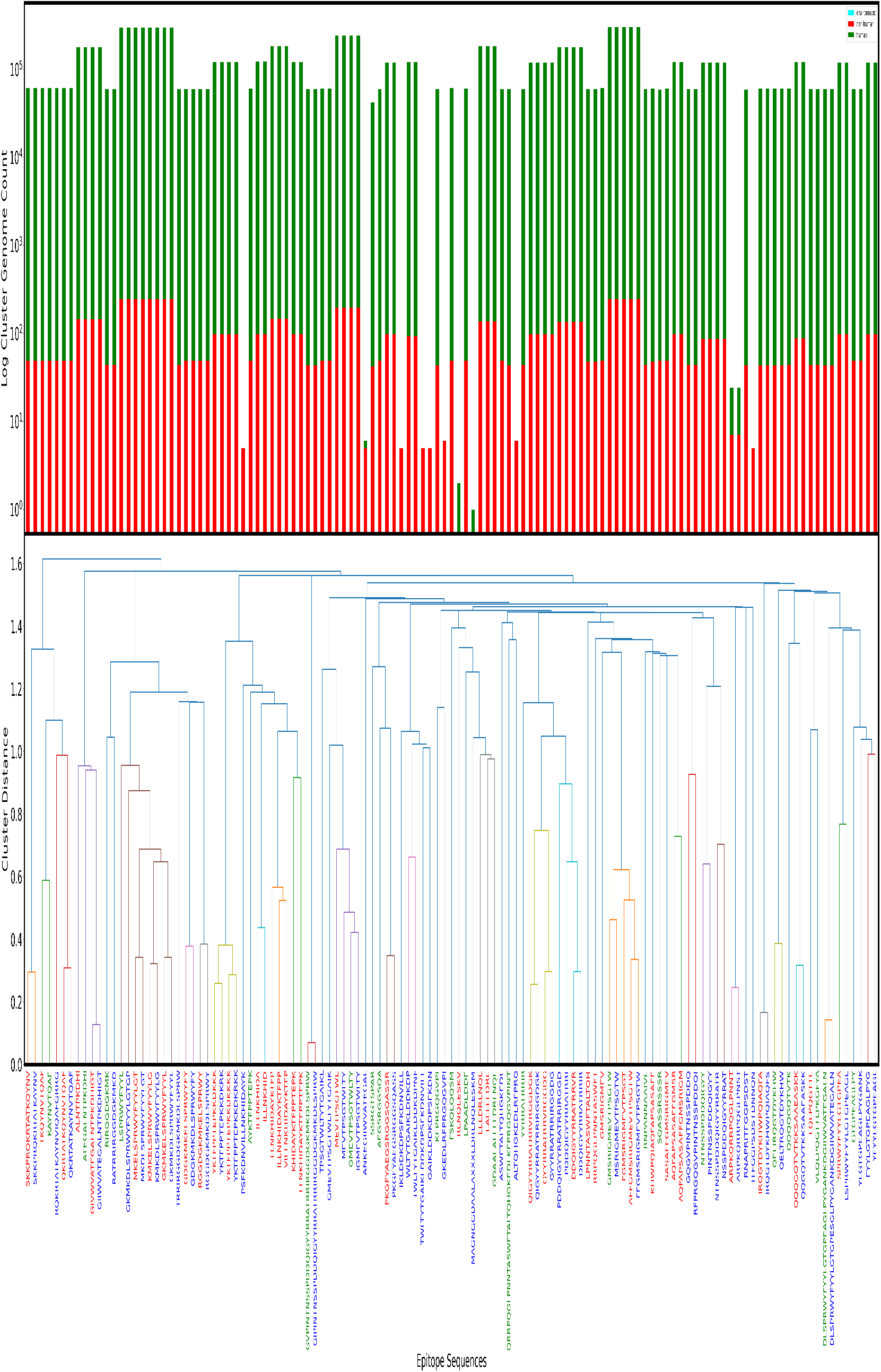
Clustering of Nucleoprotein T-Cell Epitopes *ORIGINAL* + *CANDIDATE* and their occurrence

**Figure S4.**
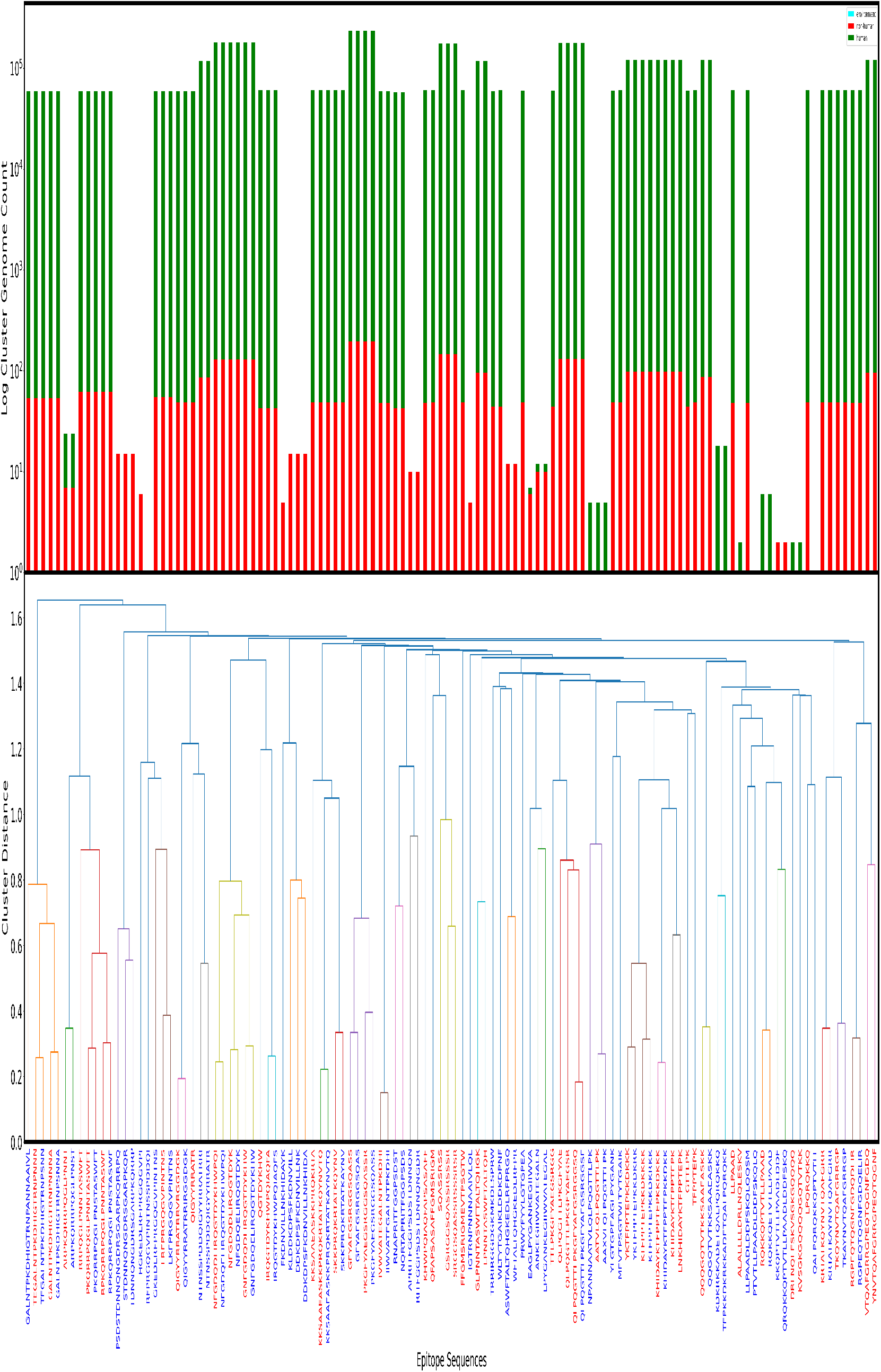
Clustering of Nucleoprotein B-Cell Epitopes *ORIGINAL* + *CANDIDATE* and their occurrence

**Figure S5.**
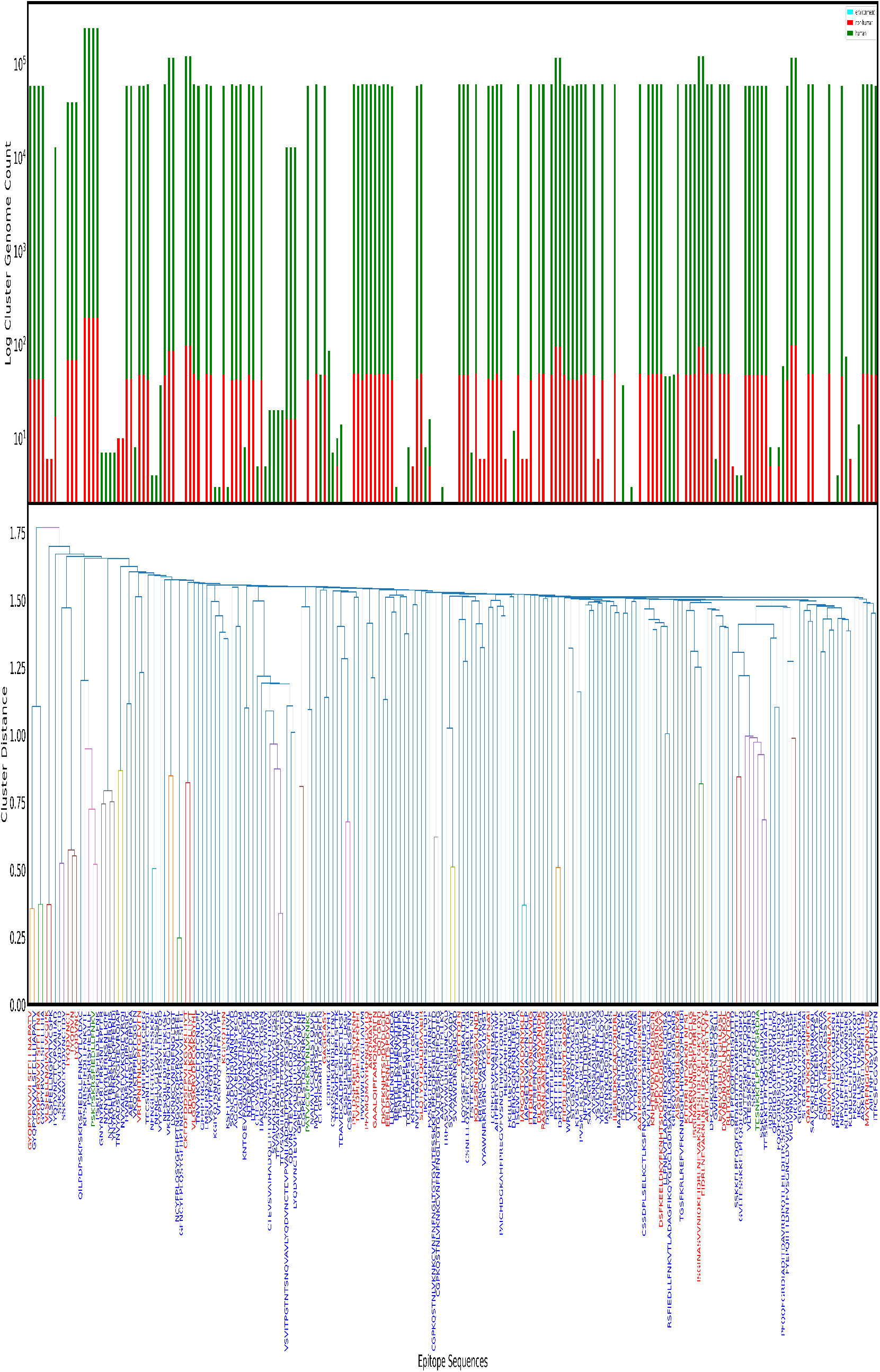
Clustering of Spike glycoprotein B-Cell Epitopes *ORIGINAL* + *CANDIDATE* and their occurrence

**Figure S6.**
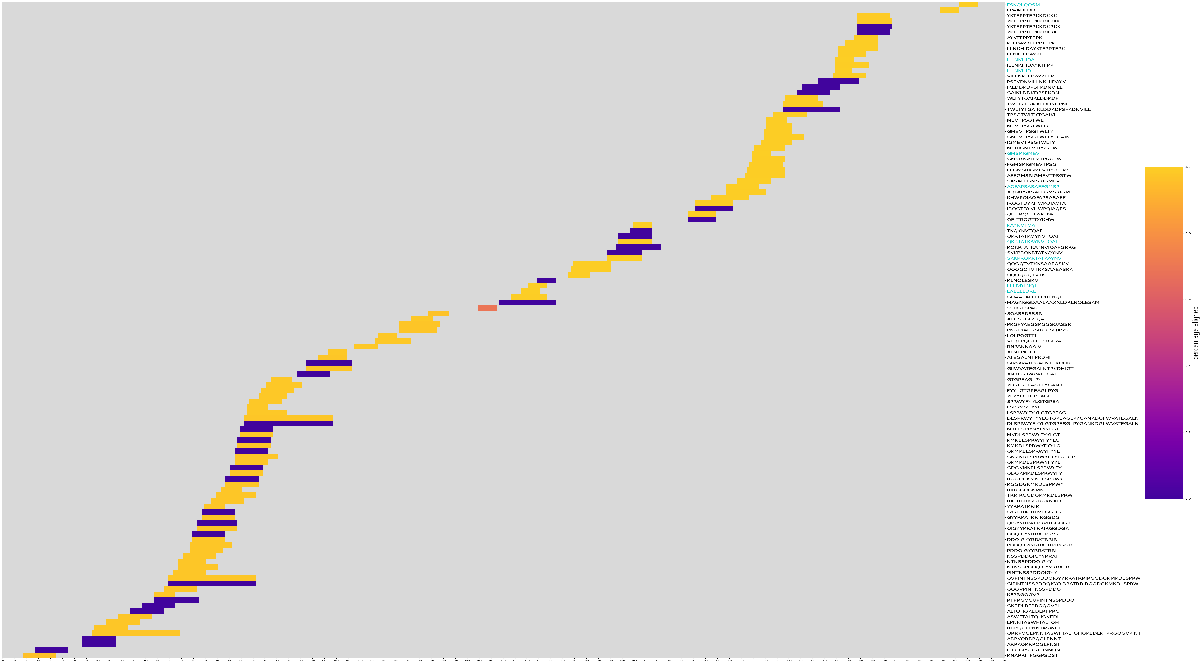
Clustermap: Position based Clustering of Nucleoprotein T-Cell Epitopes *ORIGINAL* + *CANDIDATE*

**Figure S7.**
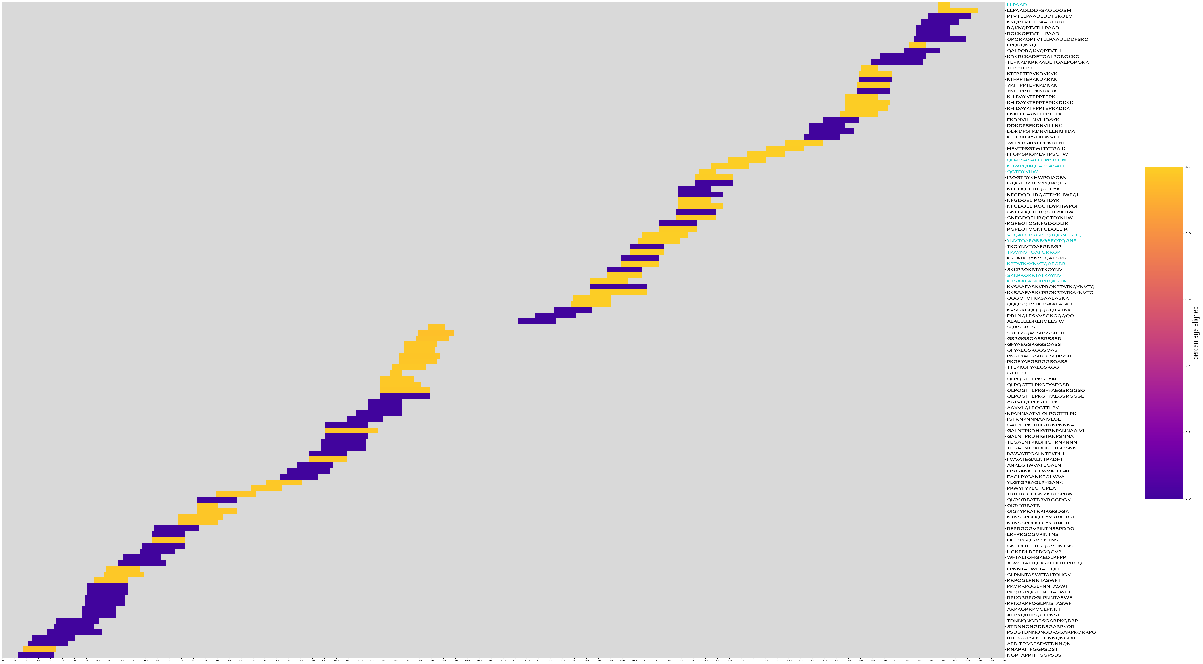
Clustermap: Position based Clustering of Nucleoprotein B-Cell Epitopes *ORIGINAL* + *CANDIDATE*

**Figure S8.**
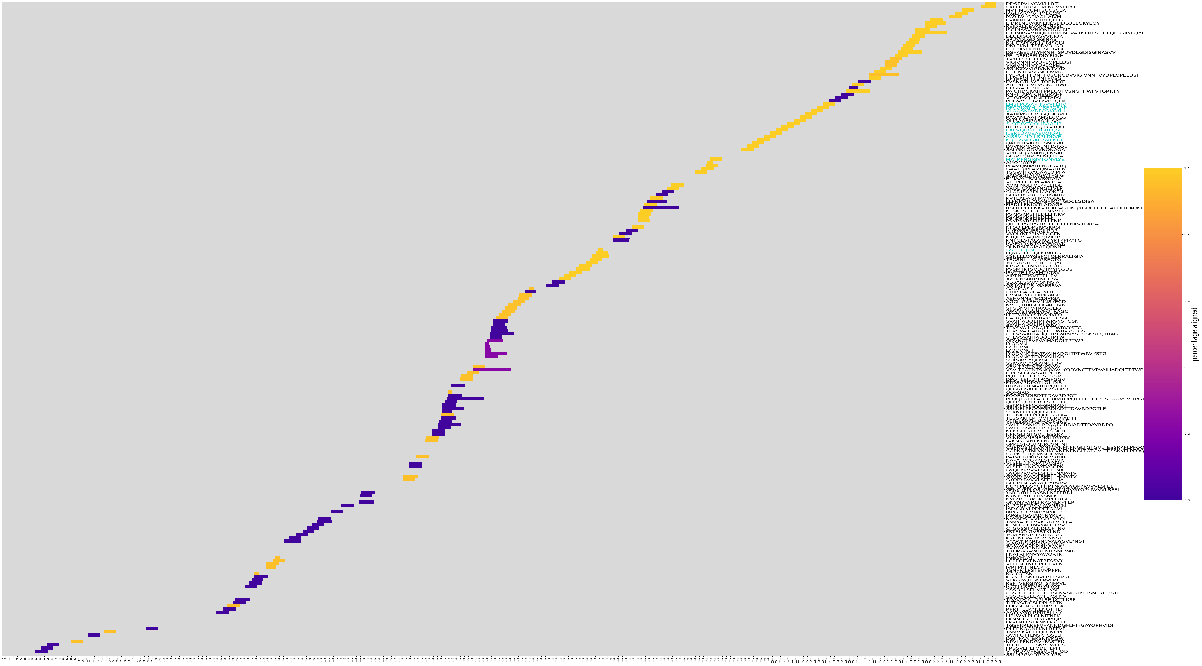
Clustermap: Position based Clustering of Spike glycoprotein B-Cell Epitopes *ORIGINAL* + *CANDIDATE*

**Figure S9.**
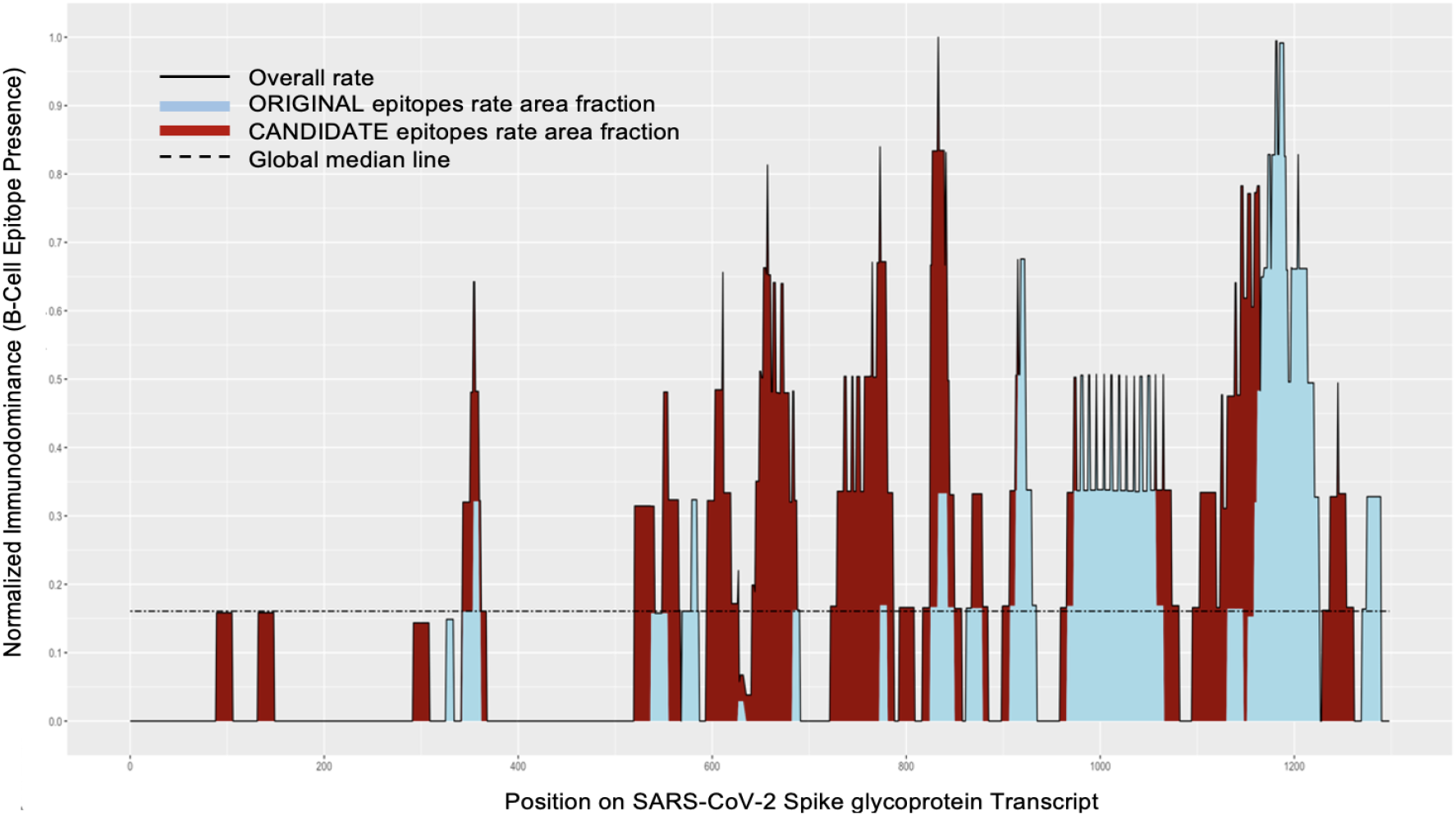
B Cell Epitope Presence on Spike Glycoprotein

**Figure S10.**
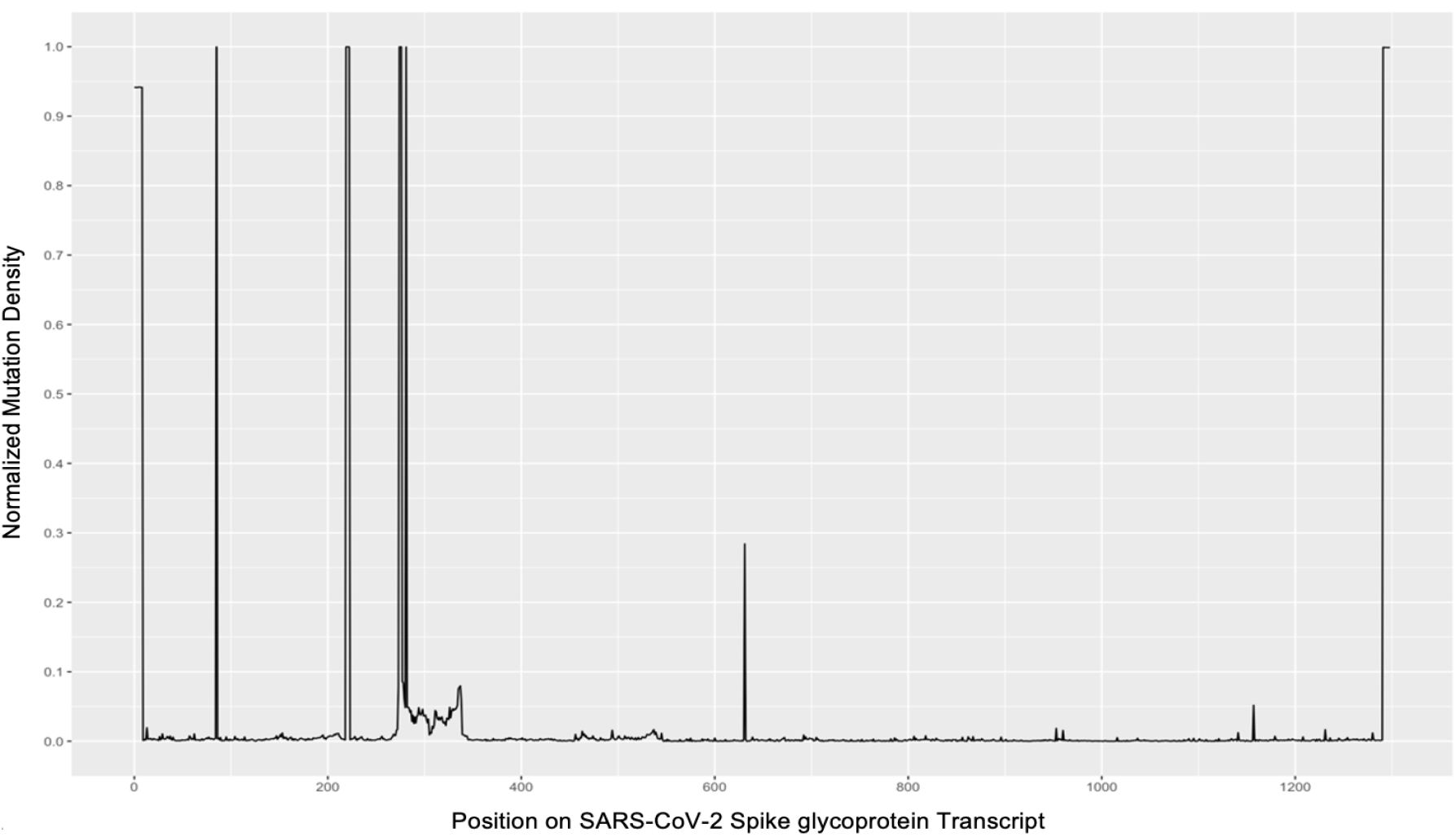
Mutation Density on Spike Glycoprotein

**Figure S11.**
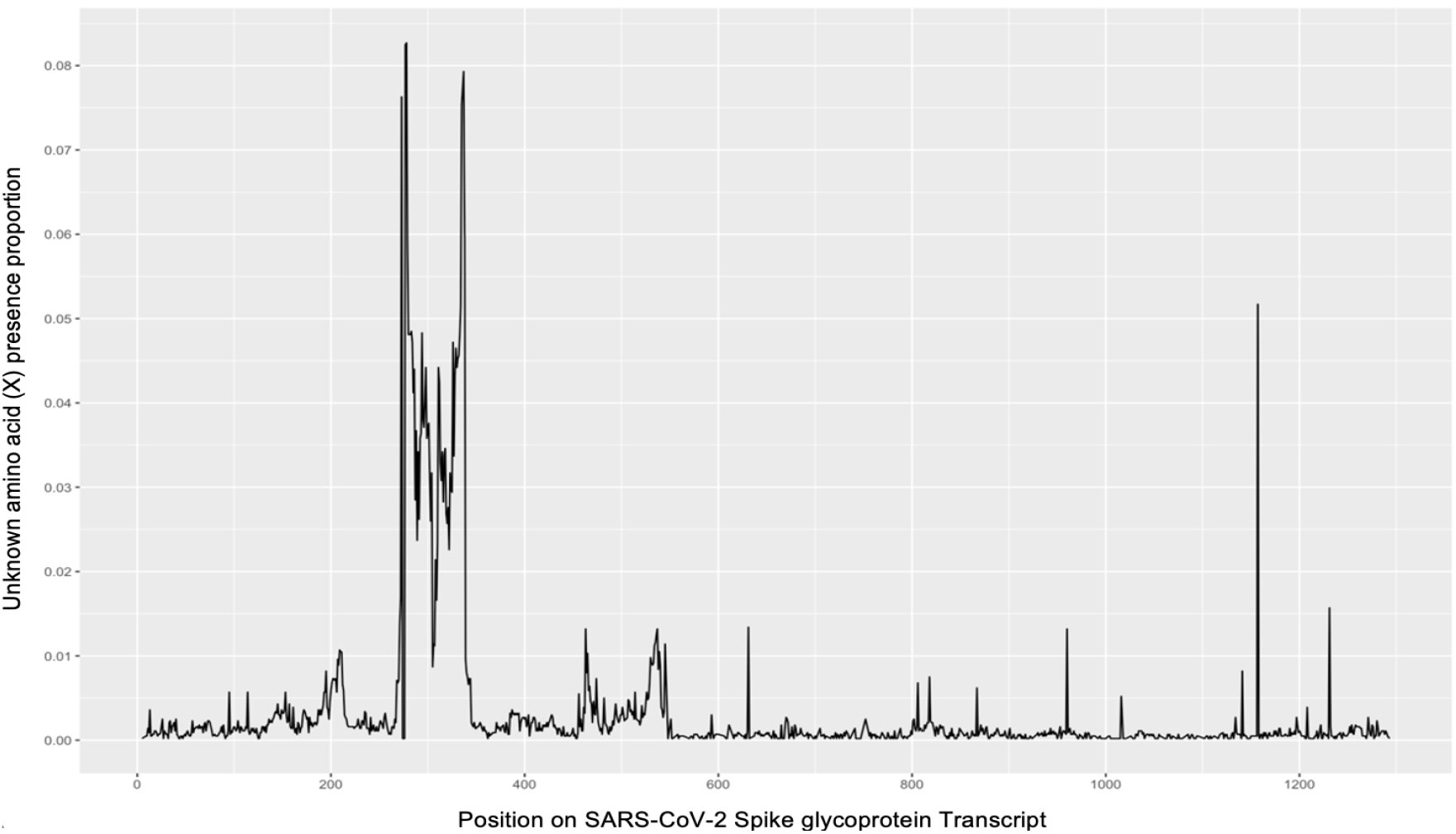
Unknown Amino Acid (X) presence proportion on Spike Glycoprotein

**Figure S12.**
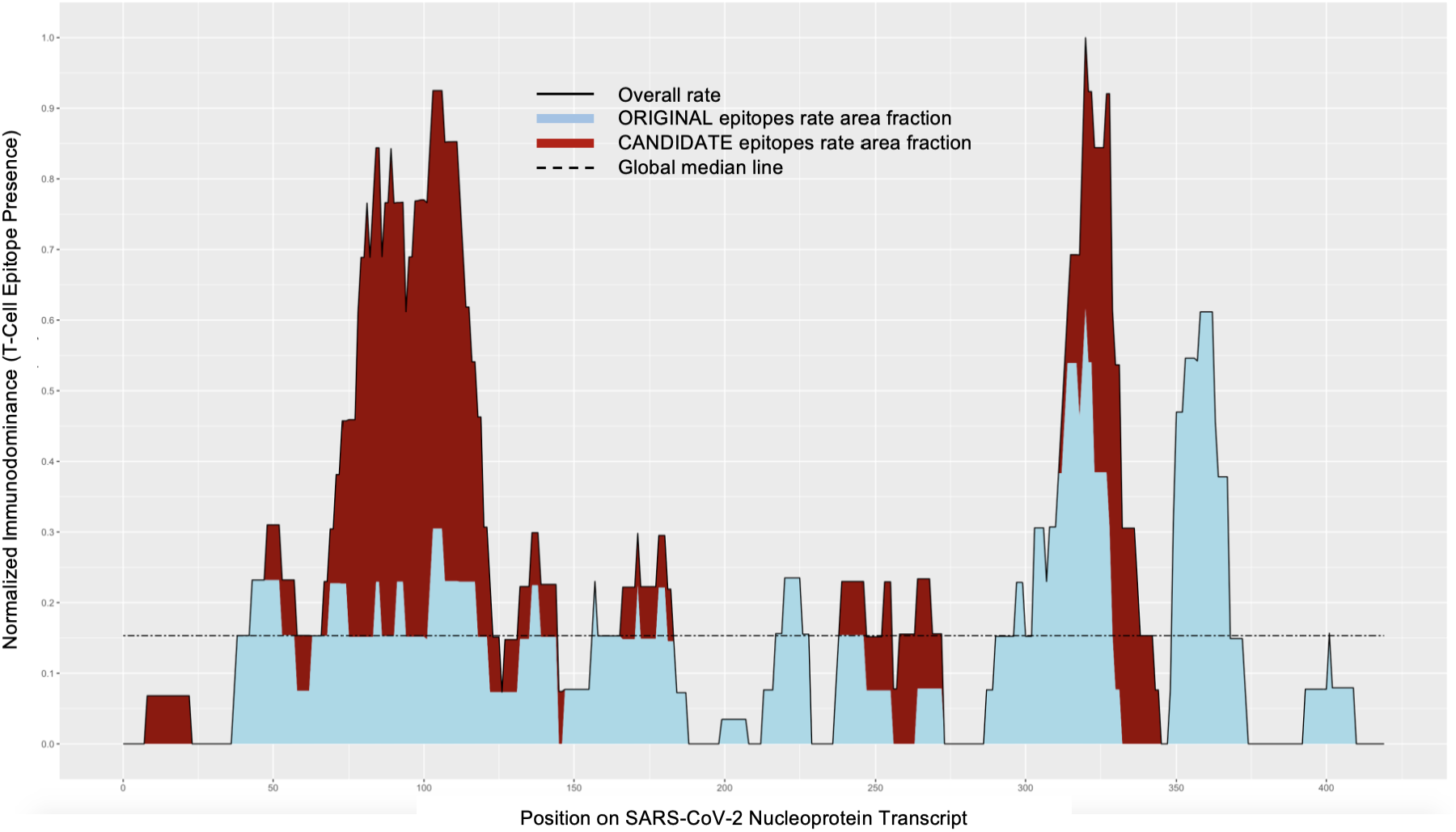
T Cell Epitope Presence on Nucleoprotein

**Figure S13.**
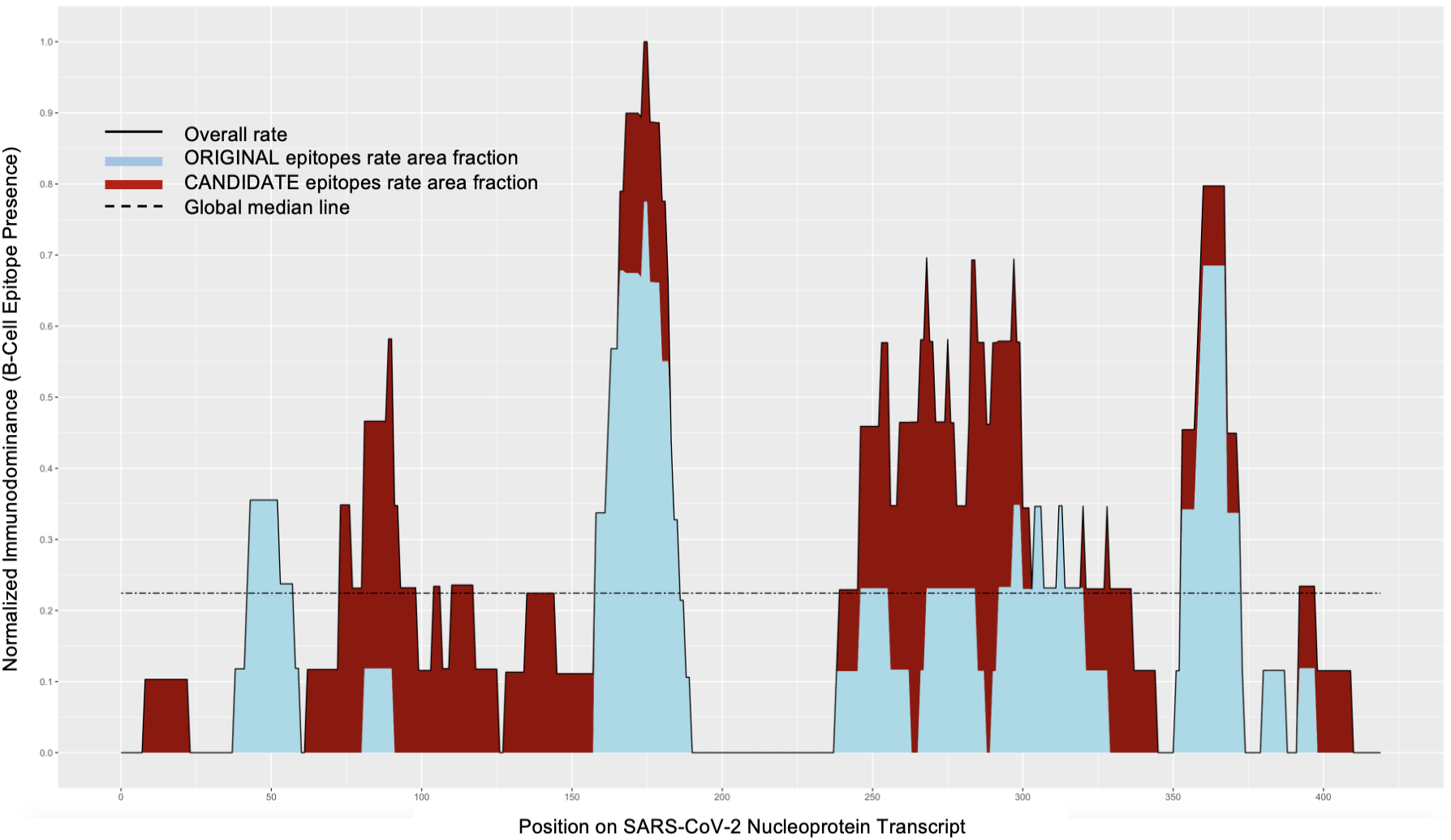
B Cell Epitope Presence on Nucleoprotein

**Figure S14.**
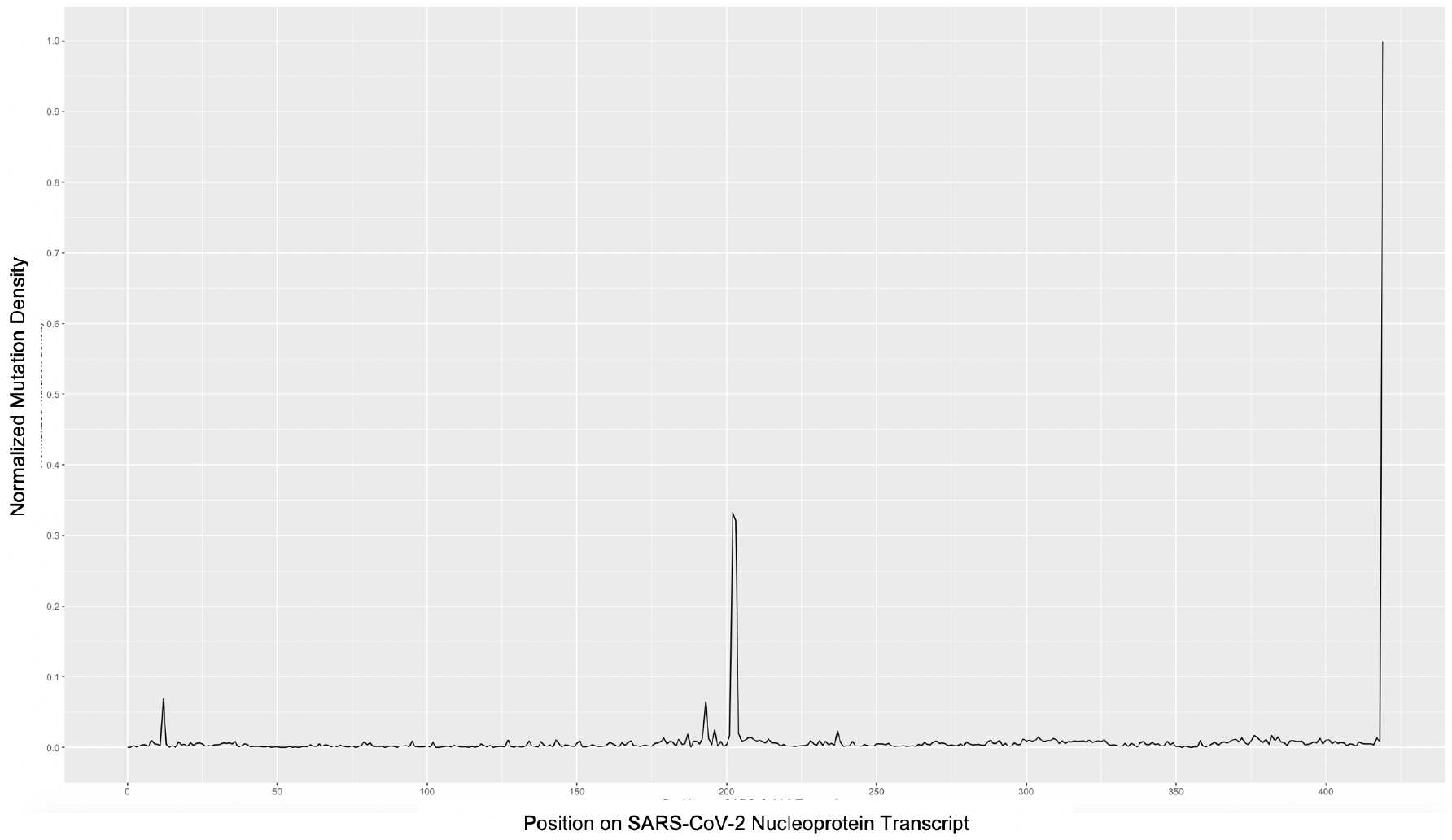
Mutation Density on Nucleoprotein

**Figure S15.**
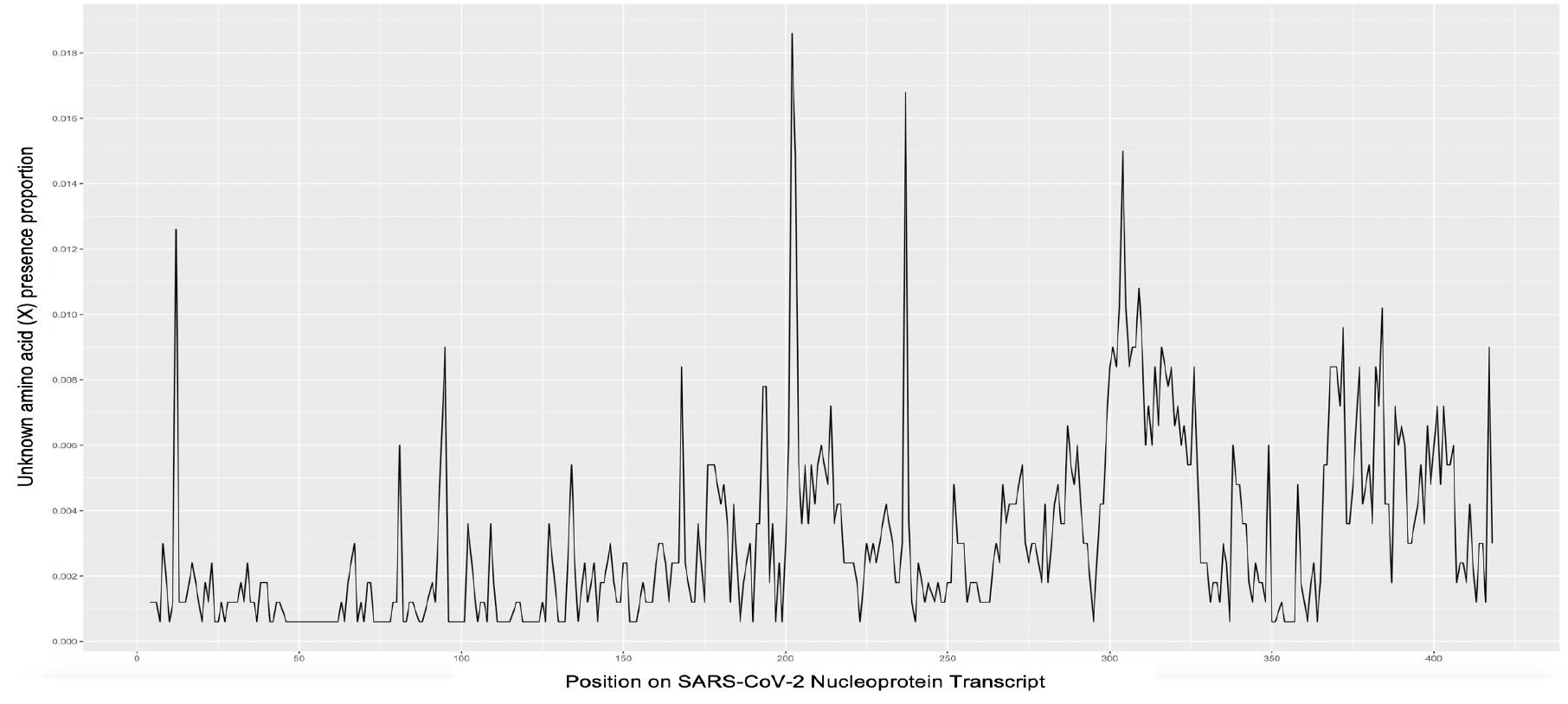
Unknown Amino Acid (X) presence proportion on Nucleoprotein

**Figure S16.**
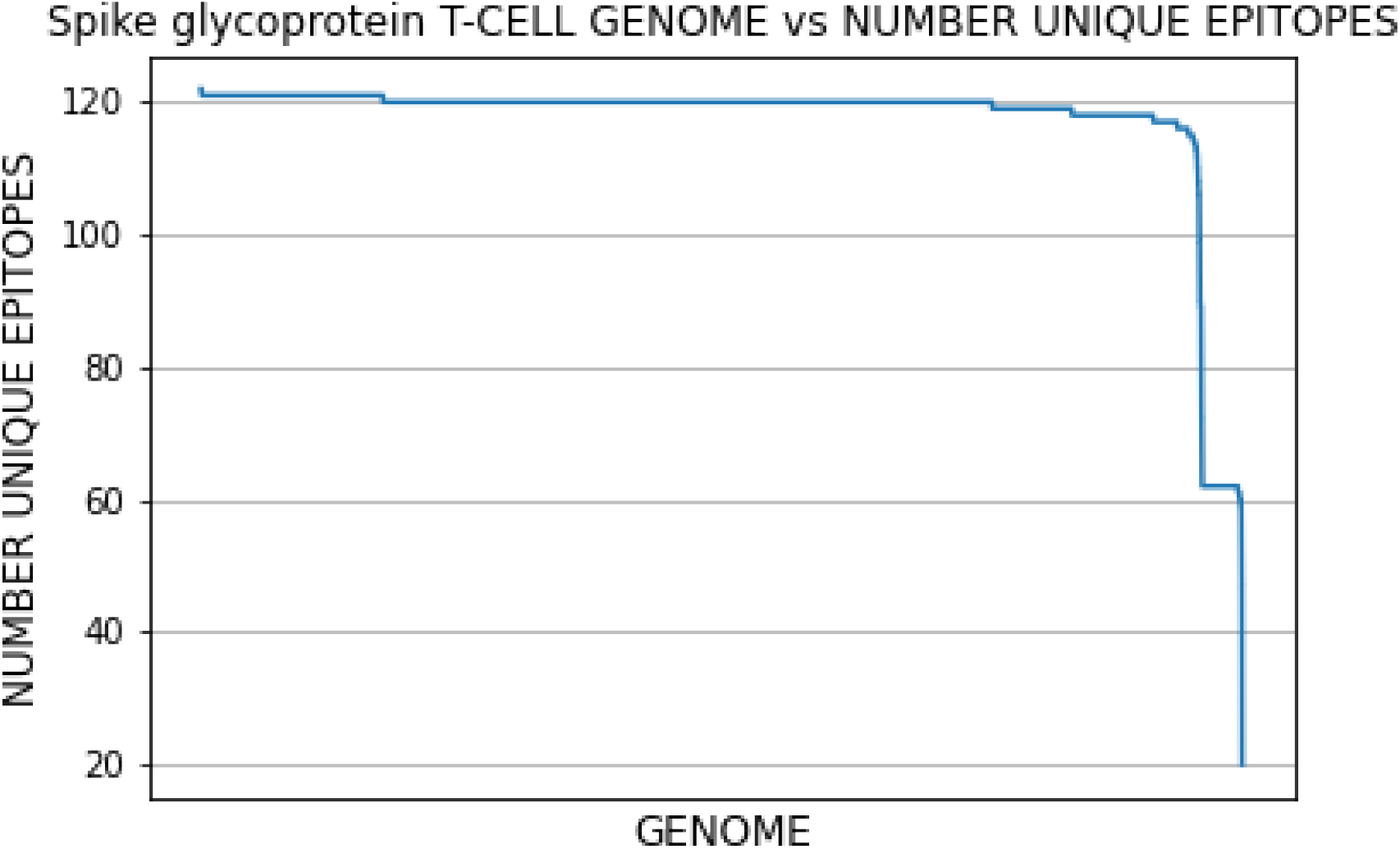
Spike glycoprotein Epitopes and SARS-CoV-2 Genomes

**Figure S17.**
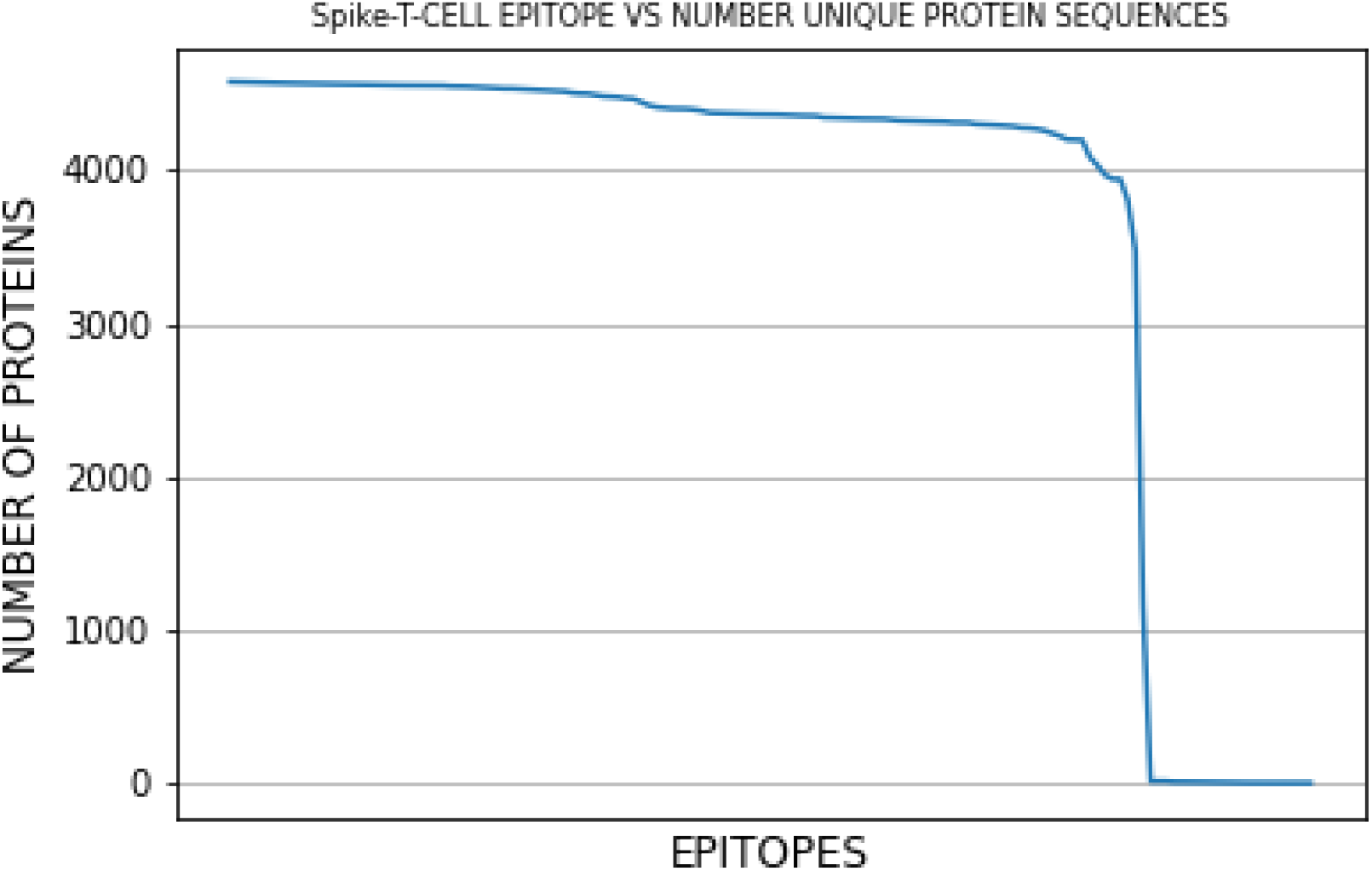
Spike Glycoprotein Epitopes and Protein Sequences

**Figure S18.**
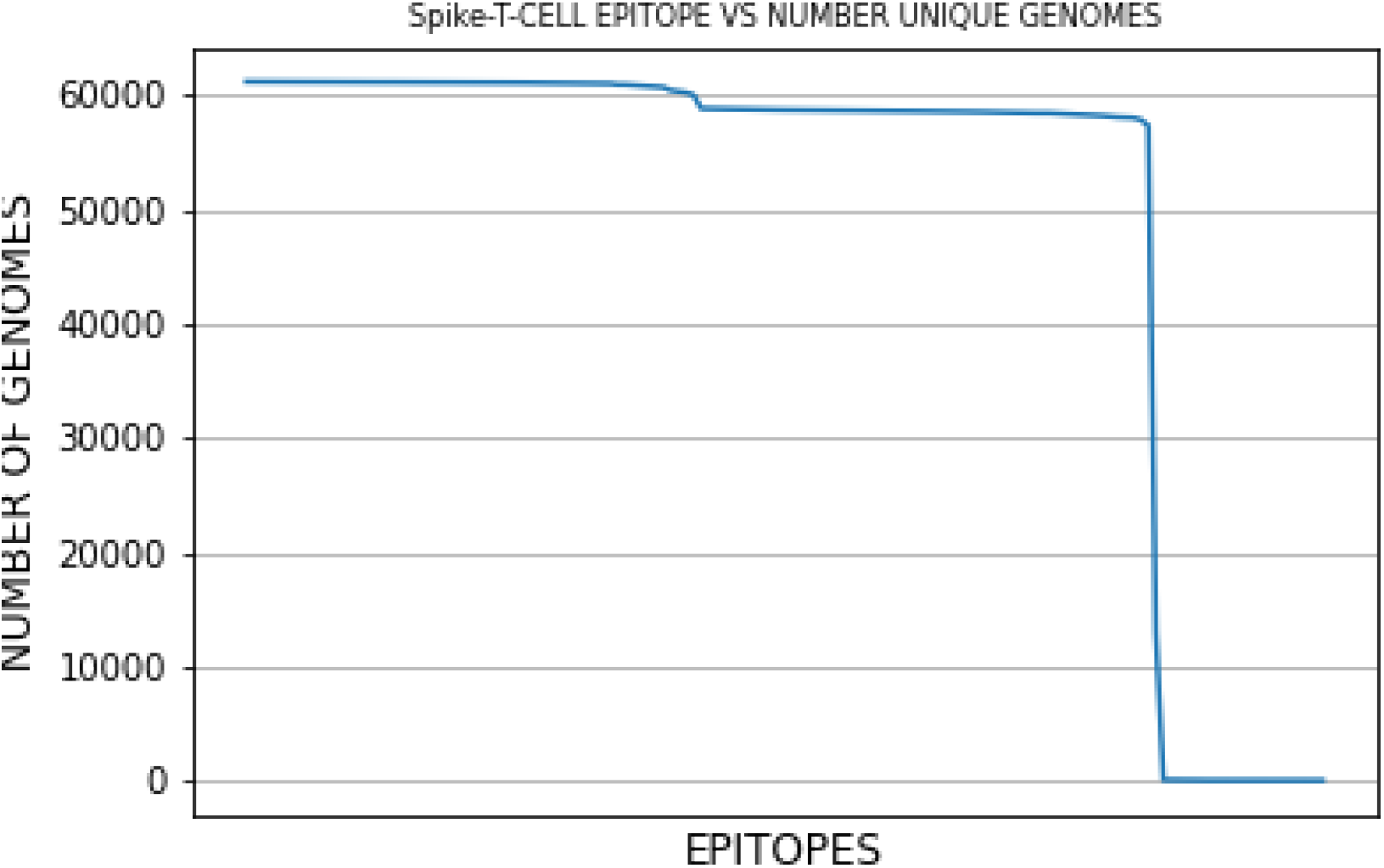
Spike Glycoprotein Epitopes and SARS-CoV-2 Genomes

**Figure S19.**
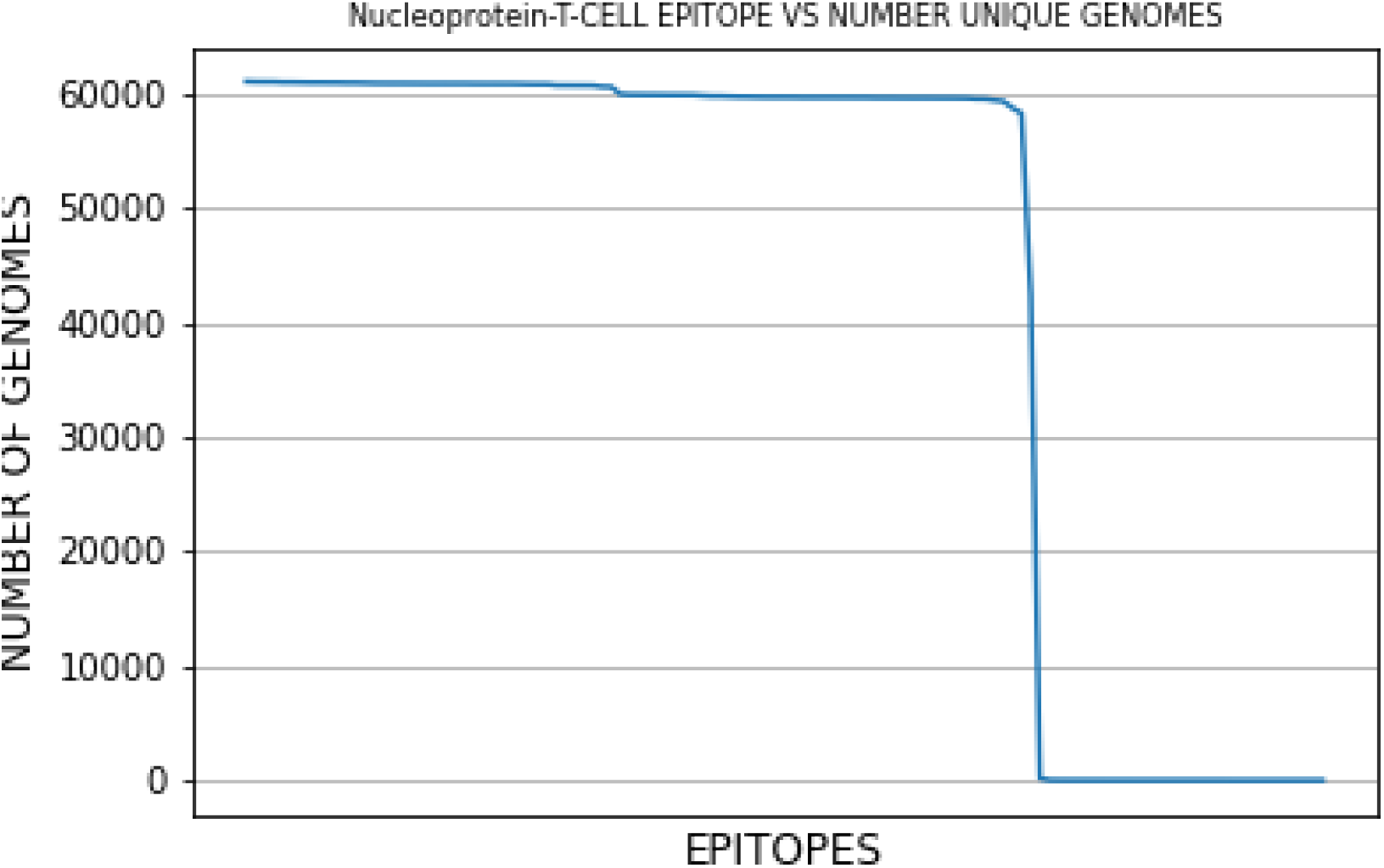
Nucleoprotein Epitopes and SARS-CoV-2 Genomes

**Figure S20.**
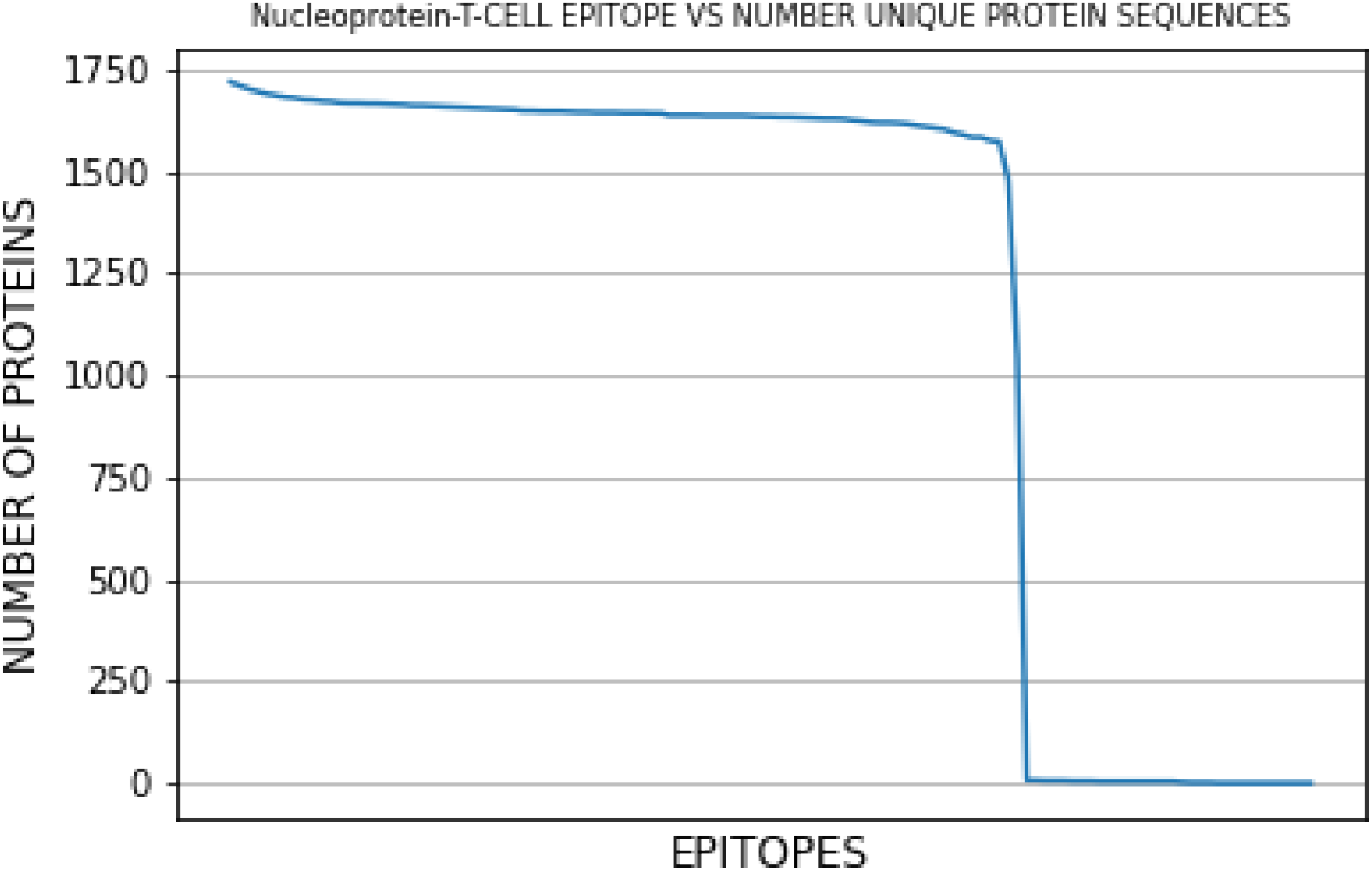
Nucleoprotein Epitopes and Protein Sequences

**Figure S21.**
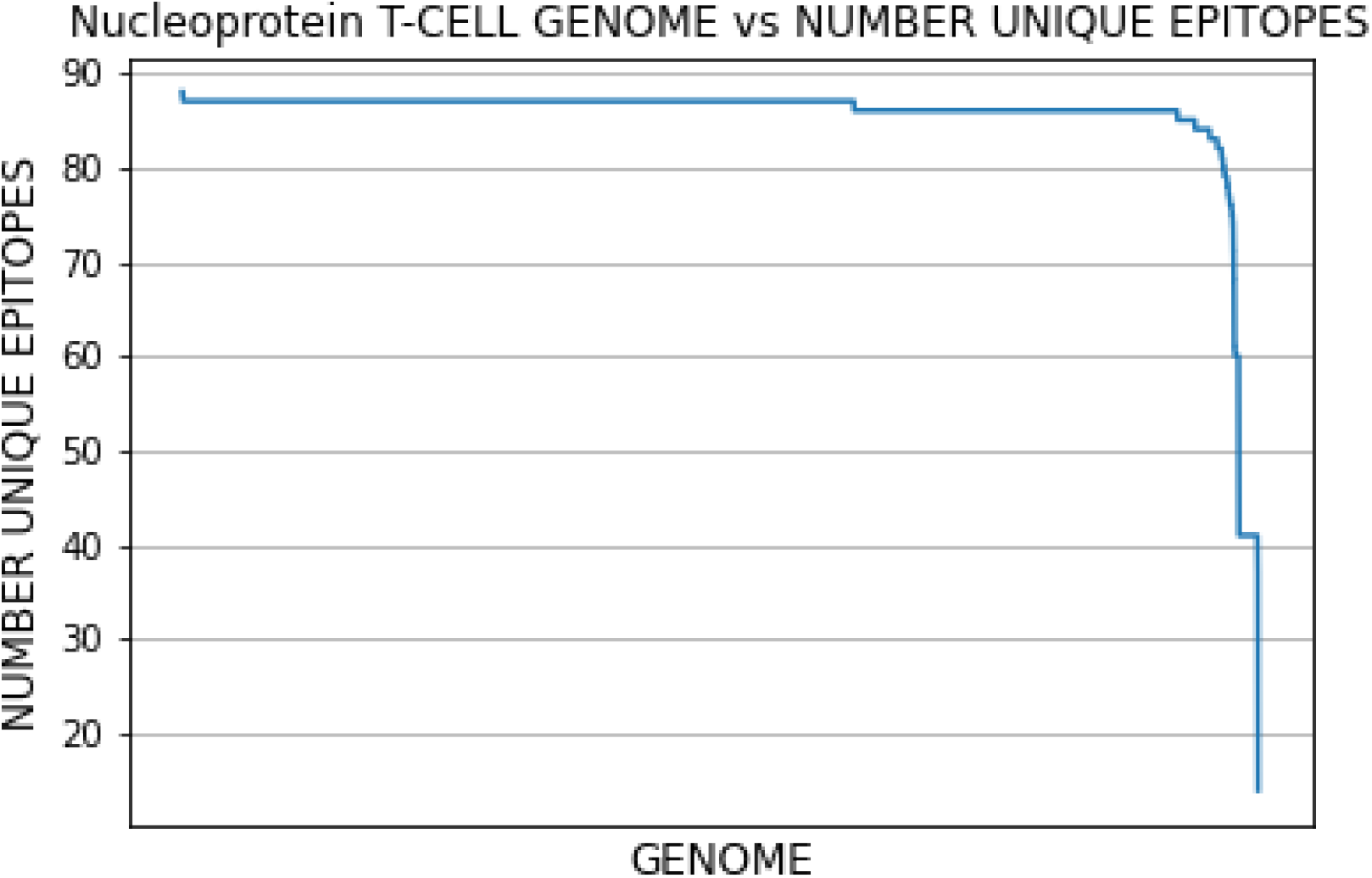
SARS-CoV-2 Genomes vs Nucleoprotein epitopes

